# Dose-dependent regulation of immune memory responses against HIV by saponin monophosphoryl lipid A nanoparticle adjuvant

**DOI:** 10.1101/2024.07.31.604373

**Authors:** Parham Ramezani-Rad, Ester Marina-Zárate, Laura Maiorino, Amber Myers, Katarzyna Kaczmarek Michaels, Ivan S Pires, Nathaniel I Bloom, Paul G Lopez, Christopher A Cottrell, Iszac Burton, Bettina Groschel, Arpan Pradhan, Gabriela Stiegler, Magdolna Budai, Daniel Kumar, Sam Pallerla, Eddy Sayeed, Sangeetha L Sagar, Sudhir Pai Kasturi, Koen K A Van Rompay, Lars Hangartner, Andreas Wagner, Dennis R Burton, William R Schief, Shane Crotty, Darrell J Irvine

## Abstract

The induction of durable protective immune responses is the main goal of prophylactic vaccines, and adjuvants play an important role as drivers of such responses. Despite advances in vaccine strategies, a safe and effective HIV vaccine remains a significant challenge. The use of an appropriate adjuvant is crucial to the success of HIV vaccines. Here we assessed the saponin/MPLA nanoparticle (SMNP) adjuvant with an HIV envelope (Env) trimer, evaluating the safety and impact of multiple variables including adjuvant dose (16-fold dose range), immunization route, and adjuvant composition on the establishment of Env-specific memory T and B cell responses (T_Mem_ and B_Mem_) and long-lived plasma cells in non-human primates. Robust B_Mem_ were detected in all groups, but a 6-fold increase was observed in the highest SMNP dose group vs. the lowest dose group. Similarly, stronger vaccine responses were induced in the highest SMNP dose for CD40L^+^OX40^+^ CD4 T_Mem_ (11-fold), IFNγ^+^ CD4 T_Mem_ (15-fold), IL21^+^ CD4 T_Mem_ (9-fold), circulating T_FH_ (3.6-fold), bone marrow plasma cells (7-fold), and binding IgG (1.3-fold). Substantial tier-2 neutralizing antibodies were only observed in the higher SMNP dose groups. These investigations highlight the dose-dependent potency of SMNP in non-human primates, which are relevant for human use and next-generation vaccines.

## Introduction

Adjuvants are substances added to vaccines that help antigens, particularly those that are weakly immunogenic, to trigger protective immune responses. Adjuvants promote activation of the innate immune system to induce adaptive immune responses (1–3). An ideal adjuvant potently facilitates broad and durable humoral and cellular adaptive immunity without triggering intolerable innate immune reactogenicity. Diverse adaptive immune responses to a vaccine may prevent or alleviate life-threating illness from exposures to the same and related pathogens. Within the adaptive immune system, T and B cells mediate targeted effector functions. Vaccine-specific CD4 and CD8 T cells are characterized by helper or cytotoxic activity, respectively. Both CD4 and CD8 T cells can develop into long-lived memory T cells (T_Mem_) capable of rapidly re-activating upon antigen re-encounter. Specific CD4 helper T cells, known as T follicular helper cells (T_FH_), are instrumental to activate B cells and facilitate B cell proliferation and differentiation during the immune reaction (4). B cells are induced to generate protective antibodies and typically affinity mature their antibodies in germinal centers (GCs). Major outputs of GC responses are memory B cells (B_Mem_) and terminally differentiated plasma cells (B_PC_). B_Mem_ serve as a resting memory population capable of re-activating upon antigen re-encounter. B_Mem_ and B_PC_, together with T_Mem_, comprise immune memory, which may have differing durability depending on the vaccine.

Emerging adjuvants are developed to induce only necessary immune effector functions(2). For example, adjuvants have been developed that are ligands for Toll-like receptors (TLRs), innate immune receptors in host cells that recognize pathogen-associated molecular patterns (PAMPs). Monophosphoryl lipid A (MPLA), a derivative of lipopolysaccharide (LPS), is a TLR4 agonist that is formulated in the licensed AS01 and AS04 adjuvants from GSK. MPLA has been shown to enhance immune responses likely via the stimulation of dendritic cells (5). Other licensed or investigated TLR agonists include TLR9 agonists (CpG DNA) or TLR7/8 agonists (3M-052), respectively, which are capable of enhancing humoral immune responses as vaccine adjuvants (6, 7).

Another important class of vaccine adjuvant compounds are saponins, triterpene glycosides originally isolated from the Chilean soapbark tree *Quillaja Saponaria*. Saponin-containing adjuvants are more potent in inducing B cell responses compared with aluminum salts (8–10) and are employed in licensed vaccines against malaria, varicella zoster virus, and SARS-CoV-2 (11–13). We recently described a novel adjuvant termed SMNP, a nanoparticle containing saponin and MPLA, which exhibited high potency in mice and non-human primates (NHPs) (8). SMNP is particularly promising for unmet vaccine needs such as HIV. No effective HIV vaccines are currently available and conventional vaccine strategies have not induced successful protection to diverse HIV strains (14, 15). Advances have been made in immunogen design and vaccine regimen including germline targeting (16–19), but all prophylactic HIV vaccine strategies depend on appropriate adjuvanticity to drive recruitment of rare B cells capable of targeting neutralizing epitopes on HIV into productive GC responses. GCs provide a highly specialized environment for B cells to affinity mature antibodies to difficult epitopes on HIV. Notably, SMNP increases lymph flow and antigen acquisition in draining lymph nodes (LNs) (8), which can augment GC responses and facilitate improved neutralizing responses (16, 20). Studies in mice provide valuable early immunological assessments for immunogens and adjuvants. However, NHPs offer a superior model for evaluating immune responses to HIV vaccines and serve as more accurate late pre-clinical models for humans due to their susceptibility to similar or identical pathogens (21). Immune memory, including durable antibody responses, is critical for the long-term benefit of prophylactic vaccines; however, in contrast to antibody responses, it is relatively uncommon to directly measure the impact of adjuvant choice and dose on B and T cell memory development in humans or NHPs, leaving significant knowledge gaps.

Accepting SMNP as a promising vaccine adjuvant, we investigated several as yet unexplored dimensions of its adjuvant activity. Saponins employed in vaccines are most often natural products. Preclinical studies to date with SMNP have employed a relatively crude saponin extract termed Quil-A, which contains a mixture of many saponin fractions (22). By contrast, for clinical adjuvant products, a highly purified saponin isolate termed QS-21 is preferred, which is composed of just two isomers of a single saponin species. QS-21 is immunologically potent and has been employed in the approved liposomal adjuvant AS01_B_ in the Shingrix™vaccine (for the prevention of shingles caused by varicella zoster) (12). The approved ISCOMs-class adjuvant Matrix-M in the Novavax COVID-19 vaccine (for the prevention of COVID-19 caused by SARS-CoV-2) employs 2 saponin fractions (Fraction-A and -C), but displays a nanoparticle structure similar to SMNP (23). Thus, key questions include: 1) How does the saponin source impact the strength and durability of immune responses? 2) How do SMNP formulations compare to a related licensed saponin-based adjuvant (AS01_B_)? 3) What are the impacts of dose or route of SMNP administration on immune responses and reactogenicity? More broadly, it is of high value to directly measure the impact of adjuvant choice and dose on B and T cell memory development in humans or NHPs. These aspects were investigated in this study to provide a broader understanding of saponin-based adjuvants in primates.

## Results

### QS-21 SMNP tested with an HIV vaccine in NHPs

We previously characterized and tested SMNP prepared using Quil-A saponin (lab-grade SMNP) in mice and NHPs (8). Clinical-grade current Good Manufacturing Practice (cGMP) SMNP synthesized using QS-21 was recently prepared for a first-in-humans clinical trial (HVTN144, ClinicalTrials.gov Identifier: NCT06033209) using a scalable manufacturing process based on tangential flow filtration (24). We collected QS-21 SMNP produced using the same process during an engineering run for this manufacturing scaleup for the present study, to assess whether scale-up manufacturing methods or the saponin source impacted responses in NHPs. Further, we aimed to carry out immunizations in rhesus macaques mirroring the planned clinical dose escalation to enable comparisons between NHP and human responses to this adjuvant. For both lab-grade SMNP prepared with Quil-A saponin and the GMP-process SMNP prepared with QS-21 saponin, the mass ratios of saponin, cholesterol, Dipalmitoylphosphatidylcholine (DPPC), and MPLA used in the adjuvant synthesis are the same (**Figure 1A**) (8). Characterization of the QS-21 SMNP showed that it had the expected cage-like nanoparticle morphology (**Figure 1B**), with particle size and composition mirroring lab-grade preparations used in our previous work (**Supplemental Table 1**) (8).

**Figure 1.**
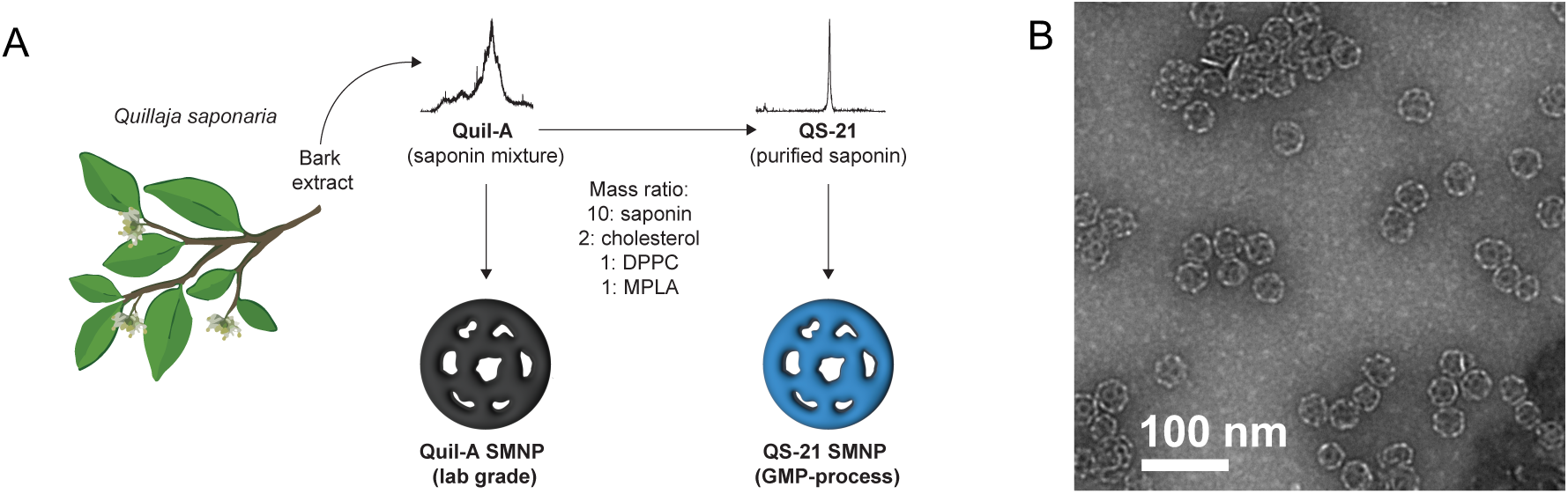
Saponin-based nanoparticles. (A) Formulation of SMNP with Quil-A (lab grade) or QS-21 (GMP-process). **(B)** Representative TEM image of QS-21 SMNP. Scale bar 100 nm.

A total of 42 rhesus macaques were split in 7 different groups with 6 animals per group. Groups 1-4 evaluated different QS-21 SMNP doses (25 µg, 50 µg, 200 µg, 400 µg). These groups employing GMP-process SMNP were compared to lab-grade Quil-A SMNP (Group 6) and AS01_B_ (Group 5). Route of vaccine immunization was also examined for QS-21 SMNP (Group 7). All groups received 100 µg HIV envelope (Env) trimer BG505 MD39 protein co-administered with the adjuvant (25). A total of 3 doses of protein plus adjuvant were administered at weeks 0, 10 and 24, unilaterally in the thighs for subcutaneous (SC) groups (Group 1-6) or intramuscularly (IM) in the deltoid (Group 7) (**Figure 2A**).

**Figure 2.**
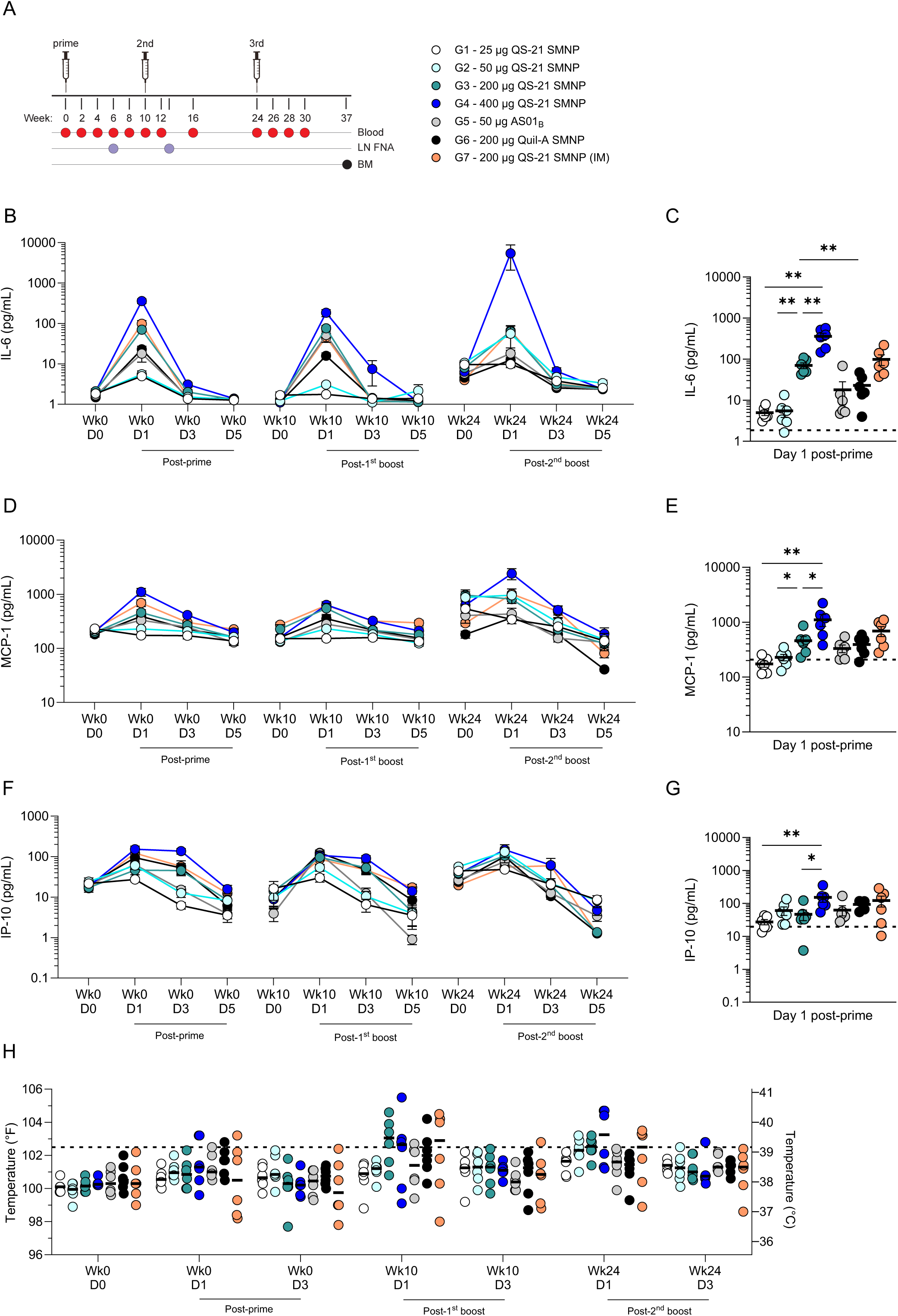
SMNP engineering run NHP study overview and reactogenicity data. (A) Overview of immunizations and sample collection schedule for NHPs. **(B-G)** Mean plasma cytokine concentration at baseline (Wk0 D0) and D1-5 following immunizations: **(B-C)** IL-6, **(D-E)** MCP-1 and **(F-G)** IP-10. **(H)** Measurement of rectal body temperature in °F (left y-axis) and °C (right y-axis) at baseline (D0) or D1-3 following immunizations. Dotted line is set at 102.5 °F/39.2 °C and represents the temperature threshold for fever in NHPs. Bars represent median per group In **(B-G)** Error bars represent mean with SEM. In **(C,E,G)** Dotted line represents the mean baseline (Wk0 D0) response of all groups. Wk= week. D= day.

After each immunization, reactogenicity and plasma cytokines were evaluated. Plasma cytokines for IL-6, MCP-1 (CCL2) and IP-10 (CXCL10) peaked at day 1 and declined to baseline levels by day 3 or 5 (**Figure 2B-G**). These cytokines are known to be involved in inflammatory responses and can promote the activity of adaptive immune responses, as well as indicate systemic reactogenicity. Somewhat surprisingly, despite covering a 16-fold dose range, a plateau in SMNP activity was not reached. At day 1 post-prime, 400 µg QS-21 SMNP (Group 4) had the highest IL-6 levels that were significantly greater than 25 µg QS-21 SMNP (Group 1 vs 4, *P* = 0.0022) and 200 µg QS-21 SMNP (Group 3 vs 4, *P* = 0.0022, **Figure 2C**). Similarly, 400 µg QS-21 SMNP (Group 4) induced higher MCP-1 levels than 25 µg QS-21 SMNP (Group 1 vs 4, *P* = 0.0022) and 200 µg QS-21 SMNP (Group 3 vs 4, *P* = 0.041; **Figure 2E**) and 400 µg QS-21 SMNP also induced greater IP-10 levels than 25 µg QS-21 SMNP (Group 1 vs 4, *P* = 0.0022) and 200 µg QS-21 SMNP (Group 3 vs 4, *P* = 0.015; **Figure 2G**). 50 µg AS01_B_ (Group 5) plasma cytokine responses were comparable to the equivalent 50 µg QS-21 SMNP dose (Group 2). Levels of IL-8, IL-10, IL-1β, IL-12p40, IL-17A, IFN-β, IL-23, TNF, IFNγ or GM-CSF did not change from baseline (data not shown). Overall, plasma cytokine concentrations were associated with QS-21 SMNP dose, but the duration of cytokine elevation did not appear to change with adjuvant dose.

Transient systemic reactogenicity (fever ≥ 102.5°F/39.2°C) was apparent in some animals, particularly with higher SMNP doses (≥ 200 µg) (**Figure 2H**). After the first immunization no substantial temperature changes were observed (**Supplemental Figure 1A**). At one day after the second immunization, fever-level temperatures were observed in a number of animals, with significant temperature elevations above baseline with 200 µg QS-21 SMNP (Group 3; **Supplemental Figure 1B**). Similar outcomes were seen after the third immunization, with significant temperature elevations with 50 µg, 200 µg and 400 µg QS-21 SMNP (Groups 2-4; **Supplemental Figure 1C**). These temperature excursions returned to baseline by day 3 in all groups. No major difference was observed between SC or IM routes of administration (Group 3 vs 7). Temperature changes with 50 µg AS01_B_ (Group 5) were overall comparable to the equivalent QS-21 SMNP dose (Group 2). Transient redness, mild rashes and swelling were observed for 1-3 days post-immunization in some animals (data not shown), but no concerning local reactogenicity events were noted.

### Dose-dependent effects of SMNP on antigen-binding memory B cells

Env-binding B_Mem_ were analyzed longitudinally by flow cytometry at different timepoints post-prime (week 10) as well as after boost 1 (weeks 12, 16 & 24) and boost 2 (weeks 26 & 30). At week 10, Env- binding B_Mem_ were detectable in all groups before a booster immunization (**Figure 3A-B**). Two weeks after the first boost (week 12), the Env-binding B_Mem_ expanded substantially in all groups (**Figure 3B-C**). The highest QS-21 SMNP dose (400 µg; Group 4) and 200 µg Quil-A SMNP (Group 6) induced the highest frequencies of Env-binding B_Mem_ responses, with median Env-binding B_Mem_ representing ∼1% of total B cells (**Figure 3C**). Env-binding B_Mem_ responses at week 12 were positively associated with QS-21 SMNP dose, with 159-fold B_Mem_ expansion in the 400 µg QS-21 SMNP group (week 12 vs 10, **Figure 3B-C**). 50 µg AS01_B_ responses (Group 5) were comparable to the corresponding 50 µg QS-21 SMNP dose (Group 2). Additionally, the 200 µg Quil-A SMNP dose group (Group 6) had similar Env-binding B_Mem_ responses compared with the equivalent dose group of QS-21 SMNP (Group 3). Delivery of 200 µg QS-21 SMNP SC (Group 3) or IM (Group 7) elicited robust and comparable Env-binding B_Mem_ responses. All Env-binding B_Mem_ frequency patterns between the groups were maintained longitudinally after the boost despite a decline observed with time in all groups (**Figure 3B, Supplemental Figure 2B**). The consistency of the decline of Env-binding B_Mem_ indicated that differentiated B_Mem_ with qualitatively similar durability were generated across the study groups (**Supplemental Figure 2D-E**), but differed in magnitude associated with the adjuvant dose. The 400 µg QS-21 SMNP (Group 4) Env-binding B_Mem_ half-life observed between weeks 12 to 16 was 9 days and over weeks 16 to 24 was 48 days. After the second boost (week 26), there was strong increase of Env-binding B_Mem_ compared to pre-boost (week 24) for all groups (**Figure 3B, D**). These Env-binding B_Mem_ responses declined similarly to what was observed with post-boost 1 responses. Overall, Env-binding B_Mem_ were induced in all QS-21 SMNP dose groups post-prime and substantially increased post-boosts. The B_Mem_ half-lives suggest a transient expansion of circulating B_Mem_ shortly after booster immunization, with the cells remaining 6 weeks later exhibiting a longer half-life.

**Figure 3.**
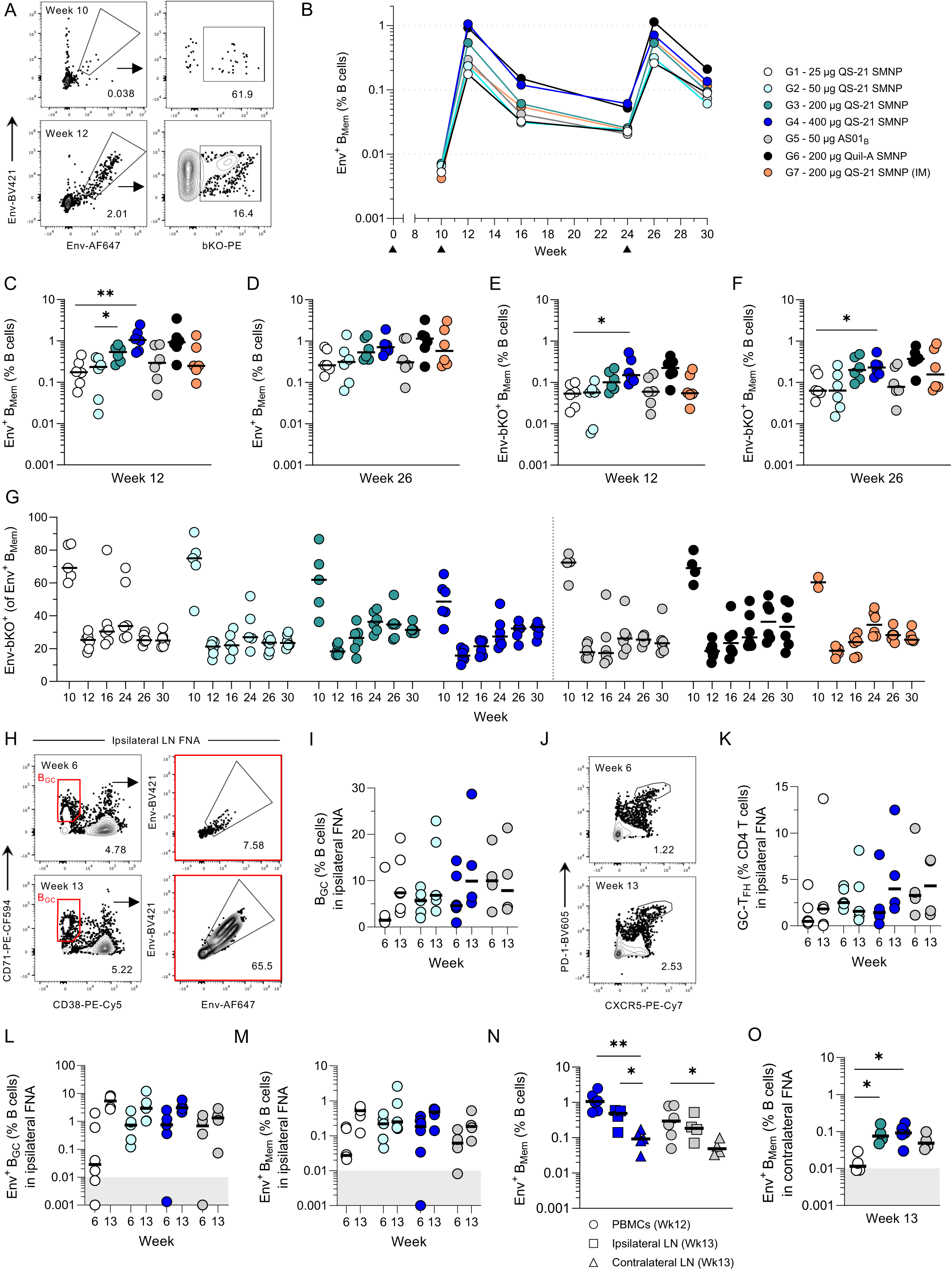
Robust longitudinal Env-binding B_Mem_ are detected with high QS-21 SMNP doses. (A) Representative flow cytometry plots of Env-binding B_Mem_ (Env-BV421^+^ Env-AF647^+^, “Env^+^”) and base knockout Env-binding B_Mem_ (bKO-PE^+^) in PBMCs at week 10 and 12. **(B)** Medians per group of Env-binding B_Mem_ as a percentage of total B cells in PBMCs at different timepoints post-immunization. Black triangles indicate times of immunizations. The limit of detection (LOD) for Env-binding B_Mem_ is 0.0006%. The immunization groups color key in panel A is used throughout Figure 3. **(C-D)** Frequency of Env-binding B_Mem_ as a percentage of B cells in PBMCs at **(C)** week 12 and **(D)** week 26. **(E-F)** Frequency of bKO-binding B_Mem_ as a percentage of B cells at **(E)** week 12 and **(F)** week 26. **(G)** Proportion of bKO-binding B_Mem_ of Env-binding B_Mem_. **(H)** Representative flow cytometry plots of total B_GC_ (CD71^+^CD38^-^) and Env-binding B_GC_ (Env-BV421^+^ Env-AF647^+^ of CD71^+^CD38^-^) in ipsilateral LN FNAs at post-prime (week 6) and post-boost (week 13). **(I)** Frequency of total B_GC_ as a percentage of B cells in ipsilateral LN FNAs at week 6 and 13. **(J)** Representative flow cytometry plots of GC-T_FH_ (PD-1^hi^CXCR5^+^) in ipsilateral LN FNAs at week 6 and 13. **(K)** Frequency of GC-T_FH_ as a percentage of CD4 T cells in ipsilateral LN FNAs at week 6 and 13. **(L)** Frequency of Env-binding B_GC_ as a percentage of B cells in ipsilateral LN FNAs at week 6 and 13. **(M)** Frequency of Env-binding B_Mem_ as a percentage of B cells in ipsilateral LN FNAs at week 6 and 13. **(N)** Frequency of Env-binding B_Mem_ as a percentage of B cells compared across 3 different tissues (PBMCs, ipsilateral LN and contralateral LN). **(O)** Frequency of Env-binding B_Mem_ as a percentage of B cells in contralateral LN FNAs at week 13. Bars represent the median per group. Grey area in **(L)**, **(M)** and **(O)** indicate the LOD. Statistical significance was determined by an unpaired two-tailed Mann-Whitney test. \**P* ≤ 0.05, \*\**P* < 0.01, \*\*\**P* < 0.001, \*\*\*\**P* < 0.0001.

Utilization of a soluble recombinant Env protein immunogen creates the issue of an immunodominant Env base surface that is otherwise buried in the membrane of HIV (26). An Env base knockout (bKO) probe (16) was used to identify B_Mem_ cells specific for non-base epitopes important for HIV neutralization (**Figure 3A**). Consistent with total Env-binding B_Mem_, bKO-binding B_Mem_ were detectable post-prime (week 10) in all groups (**Supplemental Figure 2F**). After boosting (week 12 and week 26) bKO-binding B_Mem_ were strongly enhanced and associated with the QS-21 SMNP dose (**Figure 3E-F, Supplemental Figure 2F**). The proportion of bKO-binding B_Mem_ among total Env-binders was the highest at the pre-boost timepoint (week 10) and then dropped sharply post-boost (week 12; **Figure 3G**). This outcome is consistent with the strongly competitive base responses shown before (20, 27). The proportion of bKO-binders at week 12 or later trended higher for groups receiving QS-21 or Quil-A SMNP doses ≥ 200 µg (**Figure 3G**). The bKO-binding B_Mem_ declined after the boost, but were strongly induced after the second boost (week 26) in all groups (**Supplemental Figure 2F**). Overall, durable bKO-binding B_Mem_ frequencies were induced in all groups, but were stronger in higher SMNP dose groups.

Fine needle aspiration (FNA) of LNs allows sampling of tissue adaptive immune responses, including GC responses (27, 28). Select NHP groups were analyzed for total and Env-binding B_GC_ in ipsilateral LNs post-prime (week 6) and post-boost (week 13; **Figure 3H-I**). Total B_GC_ were detected at both timepoints with comparable frequencies (**Figure 3I**). Total GC T follicular helper cells (GC-T_FH_) were also comparable across groups and timepoints (**Figure 3J-K**). Env-binding B_GC_ trended higher post-boost compared with post-prime, but were similar between 50 and 400 µg QS-21 SMNP dose groups at both timepoints (**Figure 3L**). Detection of local Env-binding B_Mem_ was apparent in the draining ipsilateral LN and trending increases were detected post-boost compared with post-prime (**Figure 3M**). The post-boost (week 13) Env-binding B_Mem_ frequencies in the ipsilateral LN were trending lower for the 400 µg QS-21 SMNP group compared with the circulating post-boost Env-binding B_Mem_ (week 12) frequencies in the blood (**Figure 3N**). For the 50 µg AS01_B_ group, week 13 Env-binding B_Mem_ frequencies in the ipsilateral LN were comparable with the circulating week 12 Env-binding B_Mem_ frequencies.

The unilateral nature of this study allowed analysis of the contralateral non-draining LN for Env- binding B cell responses. As expected, no Env-binding B_GC_ responses were detected in the contralateral LN (**Supplemental Figure 3C**). Env-binding B_Mem_ were detectable in contralateral LNs after the boost, highlighting effective re-circulation of B_Mem_ (**Figure 3O, Supplemental Figure 3C**). Env-binding B_Mem_ frequencies in contralateral LN were 8-fold higher for the 400 µg QS-21 SMNP group compared with the 25 µg QS-21 SMNP group (Group 1 vs 4, *P* = 0.016; **Figure 3O**). The frequency of Env-binding B_Mem_ in the contralateral LN was 4-5-fold lower than in the ipsilateral LN for 400 µg QS-21 SMNP (Group 4, *P* = 0.032) and trended lower for AS01_B_ (Group 5, *P* = 0.057; **Figure 3N**). Overall, the dose of QS-21 SMNP was associated with the frequency of Env-binding B_Mem_ in circulation and in LNs.

### BCR sequencing of antigen-binding memory B cells

Flow cytometric characterization of Env-binding B_Mem_ elucidated a difference in the magnitude of B_Mem_ responses based on the dose of QS-21 SMNP. To gain further insights into the quality and diversity of the Env-binding B_Mem_ at low and high adjuvant doses, single cell BCR sequencing was performed using week 12 PBMCs from all animals that received 50 µg or 400 µg QS-21 SMNP (Groups 2 and 4). Somatic hypermutation (SHM) was similar between 50 µg and 400 µg QS-21 SMNP for total Env- and bKO-binding B_Mem_ (**Figure 4A-D, Supplemental Figure 4A-D**). Regarding clonal diversity, the higher frequency of Env-binding B_Mem_ (**Figure 3C**) in the 400-µg group compared with the 50-µg group was reflected by the greater abundance of BCR clones recovered (**Figure 4E**). The clonal richness of those B_Mem_ trended higher for the higher SMNP dose (Chao1, *P* = 0.065), but did not reach statistical significance (**Figure 4F**). Similarly, the clonal abundance (rank abundance test) showed no major differences (**Figure 4G**).

**Figure 4.**
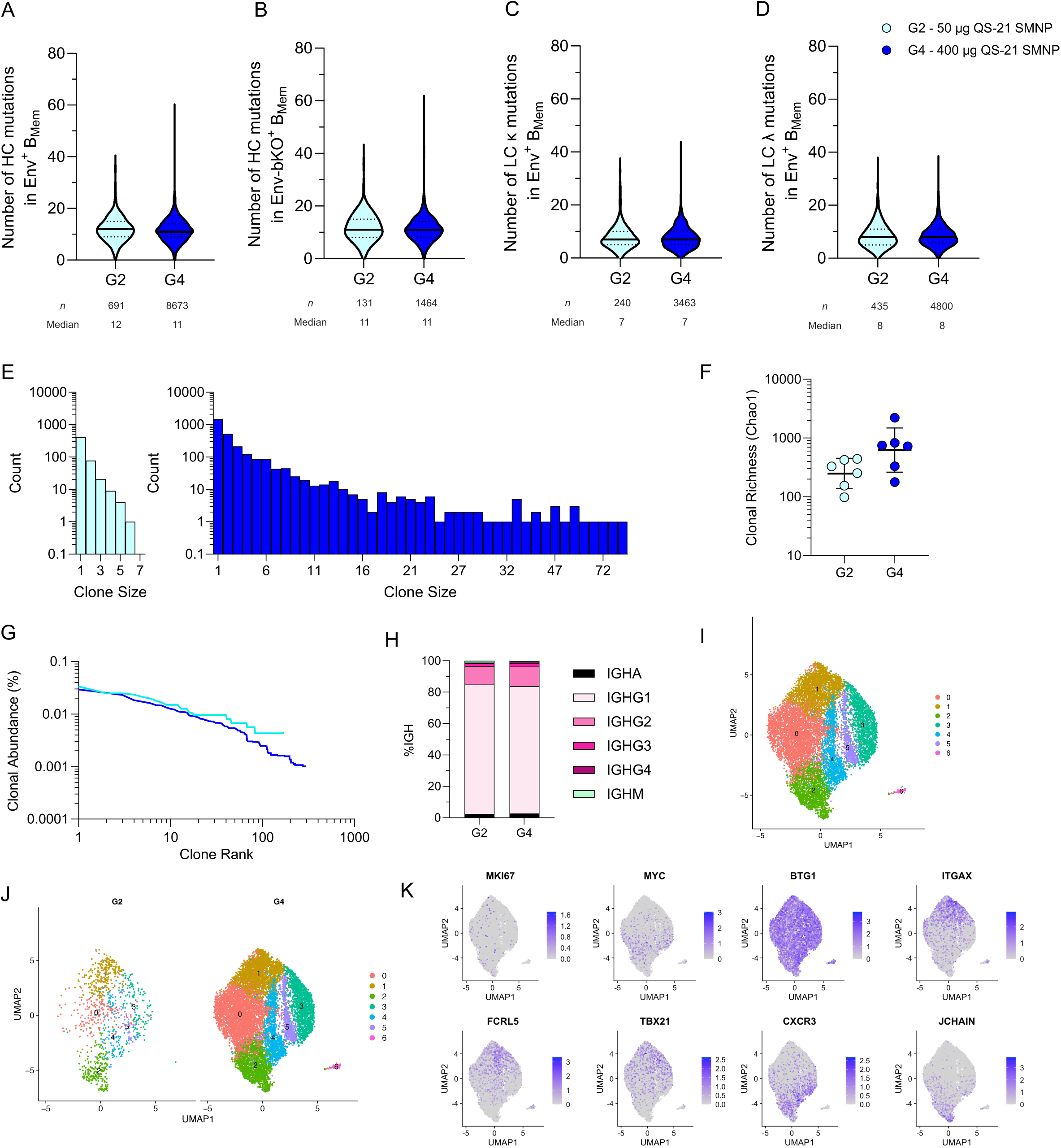
Single B cell sequencing of Env-binding B_Mem_ in low and high dose QS-21 SMNP groups. (A-B) Number of heavy chain (HC) nucleotide mutations in **(A)** Env-binding B_Mem_ and **(B)** bKO-binding B_Mem_ cells at week 12 in PBMCs. **(C-D)** Number of light chain (LC) **(C)** kappa and LC **(D)** lambda mutations in Env-binding B_Mem_ at week 12. **(E)** Distribution of clonotype sizes for Group 2 and 4. **(F)** Clonal richness (Chao1) of Env-binding B_Mem_. **(G)** Clonal abundance curve of Env-binding B_Mem_. **(H)** The proportion of immunoglobulin (Ig) isotypes of Env-binding B_Mem_. **(I-K)** Uniform manifold approximation and projection (UMAP) visualization of single-cell gene expression profiles identifying clusters of (**I**) total Env-binding B_Mem_ at week 12 (**J**) across groups and (**K**) for specific markers. **(A-D)** The solid black line and dotted lines represent the median and quartiles, respectively. The error bars in **(F)** represent the geometric mean with geometric SD. Animals were excluded from clonal abundance curve in **(G)** if less than 50 sequences were recovered. Statistical significance was determined by an unpaired two-tailed Mann-Whitney test. \**P* ≤ 0.05, \*\**P* < 0.01, \*\*\**P* < 0.001, \*\*\*\**P* < 0.0001.

Env-binding B_Mem_ were predominantly IgG1^+^ and overall Ig isotype profiles were comparable between the groups (**Figure 4H**). Single cell transcriptomic analysis of gene expression profiles of Env- binding B_Mem_ indicated major overlap in Env-binding B_Mem_ phenotypes between the two SMNP doses (**Figure 4I-K, Supplemental Figure 4E**). The proportions of Env-binding B_Mem_ were also very similar for most clusters between the groups, except some variability for a few clusters (**Supplemental Figure 4F**). The majority of cells appeared to be resting non-proliferative as observed by low levels of *MKI67* and *MYC* expression and broader expression of anti-proliferative *BTG1* (**Figure 4K**). Some atypical B_Mem_ markers including *ITGAX* (CD11c), *FCRL5* and *TBX21* (T-bet) were largely found in cluster 1. Expression of *CXCR3* appeared to be distinctly expressed compared with *ITGAX*. *CXCR3* was expressed in several clusters including cluster 2, which also contained a subcluster of cells expressing *JCHAIN,* likely indicating a small population of B_PC_ (**Figure 4K, Supplemental Figure 4G**). Overall, the diversity of B_Mem_ clonotypes and phenotypes as well as the quality of these cells was similar between low and high doses of QS-21 SMNP.

### Dose-dependent effects of SMNP on antigen-specific memory T cells

T cell responses were compared across the study groups. Acute and memory Env-specific T cell responses were investigated post-prime (week 2) and 14 weeks after boost 1 (week 24) in activation-induced marker (AIM) and intracellular cytokine staining (ICS) flow cytometric assays. Env-specific CD4 T cells responses were induced in all animals of all groups post-prime, as detected by AIM (AIM^+^ = OX40^+^CD40L^+^, **Figure 5A-B**, gated as per **Supplemental Figure 5A**). The highest dose of QS-21 SMNP (400 µg, Group 4) induced 3-fold more AIM^+^ Env-specific CD4 T cells compared to the lowest dose post-prime (Group 1 vs 4, *P* = 0.0087; **Figure 5B**). After the boost, memory Env-specific CD4 T cells were well-maintained in the two highest QS-21 SMNP doses (Group 3 & 4), but were significantly lower for the 25 and 50 µg QS-21 SMNP Groups 1 & 2 (Group 1 vs 4, *P* = 0.0022; Group 2 vs 4, *P* = 0.015; **Figure 5C**). An 11-fold difference in memory CD4 T cells was present between the lowest and highest SMNP dose (Group 1 vs 4; **Figure 5C**).

**Figure 5.**
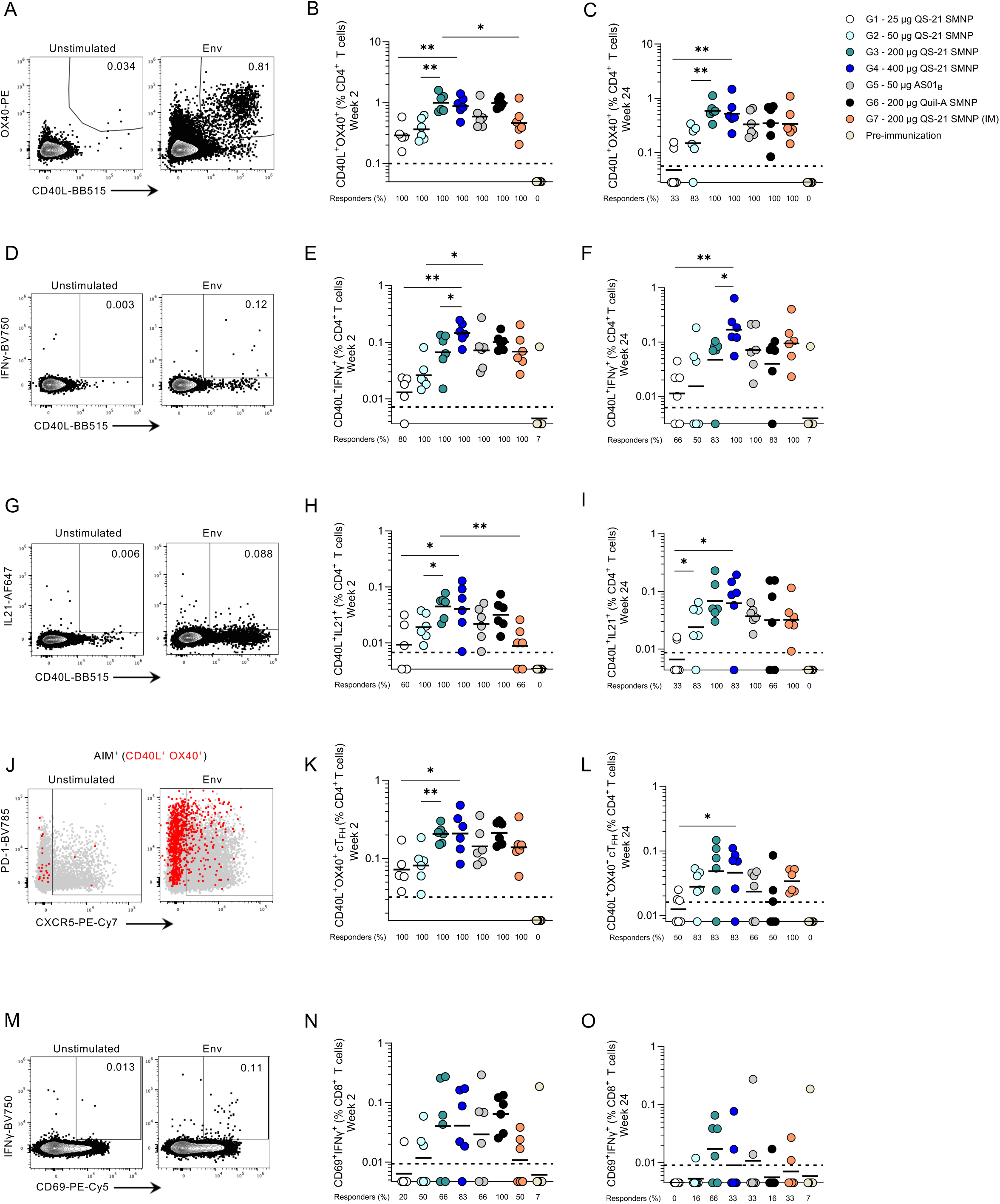
Robust Env-specific T cells are detected with all QS-21 SMNP doses post-prime. (A) Representative flow cytometry plots of AIM^+^ (OX40^+^CD40L^+^) CD4 T cells from unstimulated (negative control) and following 24-hour stimulation with a BG505 MD39 Env peptide pool at week 2. **(B-C)** Frequency of AIM^+^ (OX40^+^CD40L^+^) Env-specific CD4 T cells at **(B)** week 2 and **(C)** week 24. **(D)** Representative flow cytometry plots of CD40L^+^ IFNγ^+^ CD4 T cells from unstimulated (negative control) and following 24-hour stimulation with a BG505 MD39 Env peptide pool at week 2. **(E-F)** Frequency of CD40L^+^ IFNγ^+^ Env-specific CD4 T cells at **(E)** week 2 and **(F)** week 24. **(G)** Representative flow cytometry plots of CD40L^+^ IL-21^+^ CD4 T cells from unstimulated (negative control) and following 24-hour stimulation with a BG505 MD39 Env peptide pool at week 2. **(H-I)** Frequency of CD40L^+^ IL-21^+^ CD4 T cells at **(H)** week 2 and **(I)** week 24. **(J)** Representative flow cytometry plots of cT_FH_ (CXCR5^+^) with AIM^+^ (OX40^+^CD40L^+^) overlaid in red from unstimulated (negative control) and following 24-hour stimulation with a BG505 MD39 Env peptide pool at week 2. **(K-L)** Frequency of AIM^+^ (OX40^+^CD40L^+^) Env-specific cT_FH_ at **(K)** week 2 and **(L)** week 24. **(M)** Representative flow cytometry plots of CD69^+^IFNγ^+^ CD8 T cells from unstimulated (negative control) and following 24-hour stimulation with a BG505 MD39 Env peptide pool at week 2. **(N-O)** Frequency of CD69^+^IFNγ^+^ Env-specific CD8 T cells at **(N)** week 2 and **(O)** week 24. Bars represent the geometric mean per group. Black dotted lines indicate the limit of quantification (LOQ). Percent responders was calculated as the percent of animals above the LOQ. Graphed are the frequencies after subtracting from paired unstimulated samples. Statistical significance was determined by an unpaired two-tailed Mann-Whitney test. \**P* ≤ 0.05, \*\**P* < 0.01, \*\*\**P* < 0.001, \*\*\*\**P* < 0.0001.

QS-21 SMNP dose-dependent T cell responses were also clear in IFNγ production at acute (week 2) and memory (week 24) timepoints (Group 1 vs 4 at week 2, 11-fold, *P* = 0.0043; Group 1 vs 4 at week 24, 15-fold, *P* = 0.0022; **Figure 5D-F**, gated as per **Supplemental Figure 5A**). Similar trends were observed for Env-specific CD4 T cells producing IL-2 (Group 1 vs 4 at week 2, 3-fold, *P* = 0.0043) or TNF (Group 1 vs 4 at week 2, 8-fold, *P* = 0.0043; Group 1 vs 4 at week 24, 8-fold, *P* = 0.015; **Supplemental Figure 5F-I**). AS01_B_ (Group 5) induced higher acute CD4 T cells producing IFNγ or IL-2 than QS-21 SMNP (Group 5 vs 2, *P* =0.041 and *P* = 0.041; **Figure 5E, Supplemental Figure 5F**). IL-21 producing Env-specific CD4 T cells increased with higher QS-21 SMNP doses, with 4-fold more IL-21^+^ Env-specific CD4 T cells in Group 4 compared to Group 1 in the acute phase (*P* = 0.05; **Figure 5G-H)**, and 9-fold more IL-21^+^ memory CD4 T cells at week 24 (*P* = 0.022; **Figure 5I**).

Compared to SC, IM immunization induced 2-fold fewer acute AIM^+^ Env-specific CD4 T cells (Group 7 vs 3, *P* = 0.026; **Figure 5B**) and IL-21-producing Env-specific CD4 T cells (Group 7 vs 3, *P* = 0.0043; **Figure 5H**). Those differences were no longer significant in the memory phase (**Figure 5C, I**).

Antigen-specific circulating T_FH_ (cT_FH_) are one indicator of antigen-specific GC-T_FH_ in lymphoid organs (29–32). AS01_B_ in combination with a subunit recombinant protein immunogen has been shown to elicit higher cT_FH_ compared with a viral vector vaccination regimen (33). Acute Env-specific cT_FH_ were induced in all groups post-prime, but were highest in animals receiving 200-400 µg QS-21 SMNP or 200 µg Quil-A SMNP (Group 1 vs 4, *P* = 0.017; **Figure 5J-K**). By week 24, Env-specific memory cT_FH_ remained detectable in the majority of animals receiving 50-400 µg QS-21 SMNP (**Figure 5L**). Overall, Env-specific CD4 T cells were primed in all groups, and memory CD4 T cell frequencies were greater at higher QS- 21 SMNP doses.

CD8 T cell responses are not reported for most human or NHP adjuvanted protein vaccines. Nevertheless, multiple reports exist of CD8 T cell responses in humans and NHPs in the context of saponin-based adjuvants, including SMNP (20, 34, 35). Acute Env-specific CD8 T cell (CD69^+^IFNγ^+^) responses were elicited in some animals at week 2, with more responders at higher adjuvant dose (Group 1 vs 4, 20% to 83%; **Figure 5M-N**). Memory Env-specific CD8 T cells were only sporadically detectable at week 24 (**Figure 5O**). Overall, QS-21 SMNP induced robust and durable Env-specific CD4 T cells and some CD8 T cells, and both CD4 and CD8 T cell responses increased with higher QS-21 SMNP doses.

### SMNP dose determined bone marrow plasma cells and neutralizing antibodies

Antibodies are critical components of most vaccine immune responses. Circulating Env-binding IgG responses were longitudinally analyzed for all study groups. Env-binding IgG was detectable post-prime (week 2-10) in all groups (**Figure 6A, Supplemental Figure 6A**). Env-binding IgG substantially increased up to 13-fold post-boost (week 12 vs 10). The 200 and 400 µg QS-21 SMNP doses induced the highest amount of Env-binding IgG after the first boost (week 12 Group 1 vs 4, *P* = 0.015; Group 1 vs 3, *P* = 0.0087) and after the second boost (week 26, Group 1 vs 3, *P* = 0.041; **Figure 6B-D, Supplemental Figure 6B-D**). Importantly, Env-binding IgG decayed across the groups post-boost; however, the antibody titer stability (weeks 16-24) in the 400 µg QS-21 SMNP group was slightly better than the 25 µg QS-21 SMNP group (Group 1 vs 4, *P* = 0.0087; **Supplemental Figure 6E-F)**.

**Figure 6.**
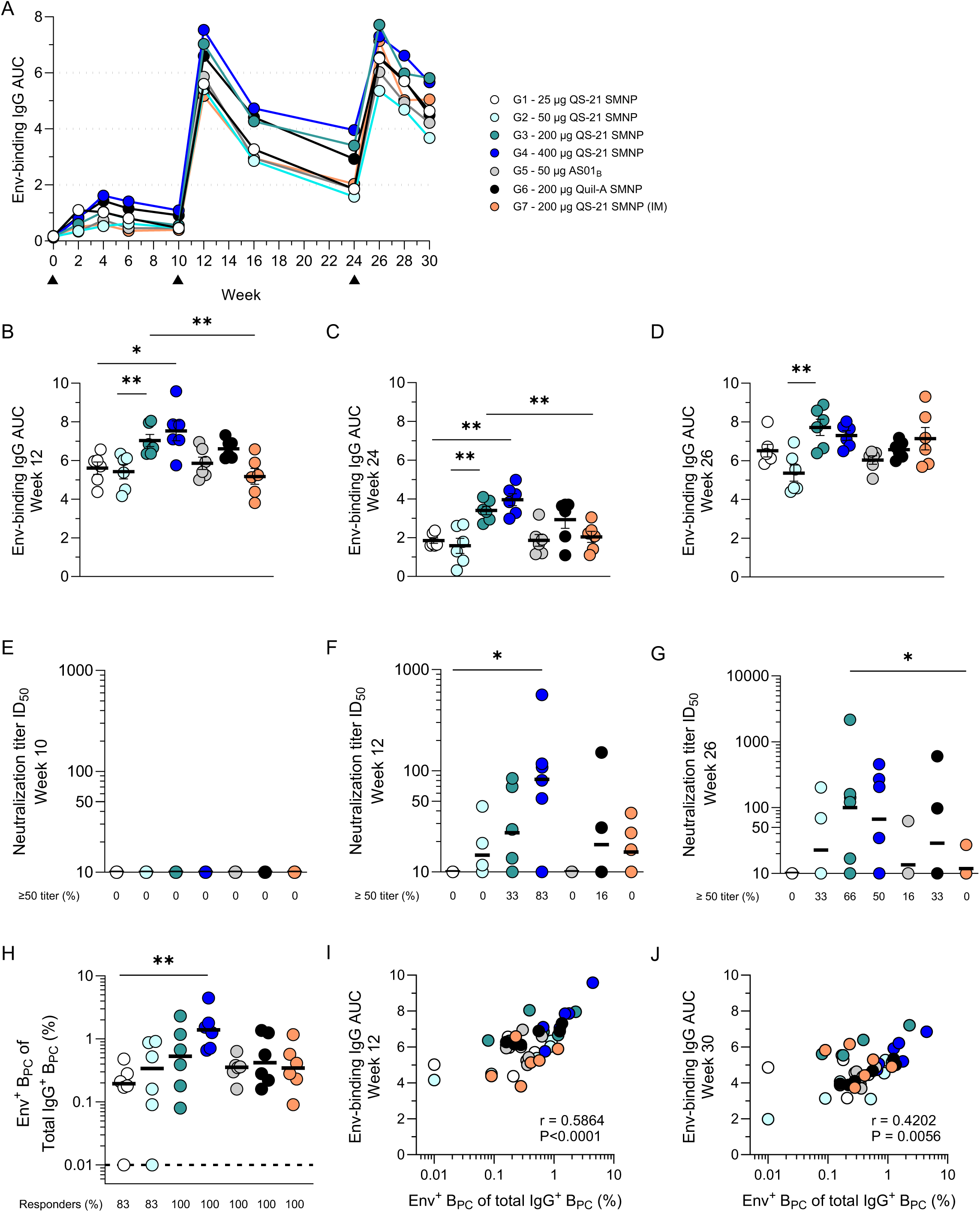
Higher induction of Env-specific BM B_PC_ and neutralizing antibodies with high QS-21 SMNP doses. (A) Mean area under the curve (AUC) of Env-binding IgG antibodies at different timepoints post-immunization. Black triangles represent the time of immunization. **(B-D)** AUC of Env-binding IgG antibodies at **(B)** week 12, **(C)** week 24 and **(D)** week 26 **(E-G)** Serum dilution at 50% inhibition of BG505 pseudovirus in a neutralization assay (ID_50_) at **(E)** week 10, **(F)** week 12 and **(G)** week 26. **(H)** Frequency of Env^+^ IgG^+^ B_PC_ as a percentage of total IgG^+^ B_PC_ at week 37. **(I-J)** Correlation between AUC of Env-binding IgG antibodies at **(I)** week 12 or **(J)** week 30 with frequency of Env^+^ IgG^+^ B_PC_. Error bars in **(B-D)** represent mean with SEM. Horizonal black lines in **(E-G)** and **(H)** indicate geometric mean and median, respectively. The dotted black line in **(H)** indicates the limit of detection (LOD) and was used to calculate percent responders. Statistical significance in **(B-H)** was determined using an unpaired two-tailed Mann-Whitney test. Data in **(I-J)** was analyzed using a Spearman’s Correlation test. \**P* ≤ 0.05, \*\**P* < 0.01, \*\*\**P* < 0.001, \*\*\*\**P* < 0.0001.

Tier-2 autologous neutralizing responses were assayed post-prime (week 10) and post-boosts (week 12 & 26). No neutralizing responses were detectable in any of the groups post-prime, as expected (**Figure 6E**). After the first boost (week 12), the highest titers were detected in the 400 µg QS-21 SMNP dose group (Group 1 vs 4, *P* = 0.015; **Figure 6F**). The 400 µg QS-21 SMNP group also had the highest amount of responding animals (∼83%, 5/6) followed by the 200 µg QS-21 SMNP group (∼67%, 4/6) and the 50 µg QS-21 SMNP group (50%, 3/6). No neutralizing antibodies were detectable in the 25 µg QS- 21 SMNP group or in the 50 µg AS01_B_ group. Compared to SC, IM immunization was inferior (week 26, Group 3 vs 7, *P* = 0.028; **Figure 6G**). Thus, SC delivery of QS-21 SMNP compared with IM mounted better polyclonal neutralizing serum antibody responses.

Bone marrow (BM) aspirates were subsequently collected to measure BM B_PC_ from all groups. Env-specific B_PC_ were detected in the majority of animals (**Figure 6H**). Interestingly, the dose-dependent trends for QS-21 SMNP were consistent with other immune responses, with the 400–µg group inducing the highest amount of Env-specific B_PC_ (Group 1 vs 4, *P* = 0.0022; **Figure 6H**). BM B_PC_ frequencies correlated significantly with circulating Env-binding IgG titers (**Figure 6I-J**). Overall, 400 µg QS-21 SMNP induced the highest amount of Env-binding IgG, neutralizing antibodies, and BM B_PC_ responses.

## Discussion

This study shows that QS-21 SMNP is a potent adjuvant for recombinant protein immunizations, with a promising safety and reactogenicity profile in NHPs. The SMNP dose regulates the magnitude and quality of the immune memory responses. The lowest tested dose of QS-21 SMNP at 25 µg in this study was capable of inducing immune memory. Doses of up to 400 µg QS-21 SMNP were well tolerated and induced the strongest responses, including neutralizing antibodies and antigen-specific B_Mem_, B_PC_ and T_Mem_. The comparison to AS01_B_ is an important clinically relevant comparator for QS-21 SMNP. 50 µg of QS-21 SMNP or AS01_B_ elicited comparable immune responses. This is notable given that SMNP contains 10-fold less MPLA than AS01_B_, and it is plausible that SMNP plus protein can be co-formulated, as ISCOMs-related adjuvants in clinical use are (36), unlike AS01_B_, which cannot be stored premixed with protein. Overall, QS-21 SMNP is a powerful and potent new adjuvant for immune responses associated with protective immunity.

Durability of immune responses is critical to ensure ongoing vaccine protection in the absence of repeated boosters. The data reported herein highlight that SMNP has the ability to induce durable B and T cell memory. Several lines of evidence indicate better immune memory with higher QS-21 SMNP doses. Env-specific CD4 T_Mem_ and B_PC_ were more robustly and reliably detected in the 400 µg QS-21 SMNP animals compared with the 25 µg QS-21 SMNP group. Furthermore, Env-binding IgG responses had less decay post-boost (weeks 16-24) in the 400 µg QS-21 SMNP group indicating more durable circulating antibody responses. The strong correlation between Env-binding B_PC_ and Env-binding IgG further supported these findings. Env-binding B_Mem_ frequencies decayed with similar kinetics in high and low dose QS-21 SMNP groups post-boost, but the 400 µg QS-21 SMNP group had higher frequencies compared with the 25 µg QS-21 SMNP at week 24. These results indicate that SMNP induces dose-dependent Env-binding B_Mem_ responses that last longer post-immunization.

While the magnitude of Env-binding B_Mem_ was higher with 400 µg QS-21 SMNP compared with 50 µg QS-21 SMNP after the booster immunization, single B cell sequencing revealed similar qualities early after the boost in both cases. Based on the similarities of Env-binding B_Mem_ between different doses of QS-21 SMNP, it is plausible that their affinities were generally comparable as well. Antigen-specific B_Mem_ can increase their SHM after two immunizations in NHPs and humans (37–39). This phenomenon may predominantly occur through recall responses of B_Mem_ into GCs (16, 40). The 50 and 400 µg QS-21 SMNP groups may have different GC longevities, which could impact the level of recall responses and thereby the diversity and quality of Env-binding B_Mem_ at later timepoints. The induction of more abundant B_Mem_ clonotypes in the 400 µg QS-21 SMNP group compared with the 50-µg group may also increase the chance of recruiting rare clones with neutralizing potential into the response. Overall, vaccine strategies capable of inducing durable immune responses will be facilitated by in-depth understanding of how different antigen-specific B and T cell compartments are induced and maintained. Protective vaccine responses generally require the induction of robust antigen-specific T cell activity. AS01 adjuvants, including AS01_B_, were selected for advancement to clinic based on their ability to induce enhanced cellular immunity in addition to humoral immunity (41). T cell response induction by SMNP was of particular interest for this study. Env-specific CD4 T cells were primed at all SMNP doses in this study. Consistently high CD4 T cell responses were elicited with SMNP doses ≥ 50 µg. Env-specific memory CD4 T cells were generated at all SMNP doses. Env-specific memory cT_FH_ and CD4 T cells producing IFNγ or IL-21 were more abundant with higher QS-21 SMNP doses. Overall, SMNP at the doses tested was capable of inducing substantial frequencies of antigen-specific CD4 T cells, with robust functionalities, indicative of substantial capacity to aid with vaccine antibody responses and potentially protective immunity.

Env-specific CD8 T cells were also detected in this study. Env-specific CD8 T cells varied among the study groups, but were primed more consistently with SC delivery of QS-21 or Quil-A SMNP doses ≥ 200 µg. More work is required to determine how protein immunogen with adjuvants prime antigen-specific CD8 T cells and what specific role these cells may play in protective immunity.

We administered immunizations unilaterally in this study to model the most common situation in human vaccination. The neutralizing responses after unilateral administrations here for the highest SMNP dose group (400 µg QS-21 SMNP) were slightly lower than previously published bilateral administration of 375 µg (187.5 µg/side) or 750 µg (375 µg/side) Quil-A SMNP with equivalent doses of BG505 MD39 (8, 20). Theoretically, bilateral immunizations can reach twice as much LN engagement than unilateral, thus increasing the odds of recruiting rare clones with neutralizing capabilities. Although the dose and saponin source are not identical in these studies, the outcomes indicate it is plausible that bilateral immunization increases neutralizing antibody responses. However, this needs to be directly tested in future studies. SC administrations showed stronger responses than IM, including neutralizing antibody responses. SC drainage is less restricted than IM (42, 43), which can increase LN engagement. Although the anatomical site (thigh vs. arm) differs here, this finding is consistent with previous reports showing lower tier-2 autologous neutralizing response via IM than SC delivery in the thighs of NHPs (42). In sum, SC delivery of QS-21 SMNP compared with IM delivery led to stronger acute CD4 T cell responses and neutralizing antibody responses.

Recent advances in QS-21 production using tobacco plants or yeast as a heterologous expression system instead of the Chilean soapbark tree are encouraging to provide sustainable sources for this rare and crucial molecule (44, 45). Although we show QS-21 is an excellent saponin in terms of potency and safety to formulate in SMNP, the proven safety of Quil-A in formulations with phospholipids and cholesterol in animal models should not be neglected. This potentially suggests other more abundant saponins could be considered for their applications in vaccine adjuvants as well. It would be worthwhile to compare different saponin fractions in SMNP formulation to investigate the potential different roles of QS-21 and other fractions on immune responses and their linked mechanism. Initial innate immune responses are likely important in the optimal induction of these durable adaptive immune responses. A better understanding of the mechanism of action of adjuvants will be important to build an arsenal of different immune activators to generate durable responses.

In sum, these results are promising and encouraging for the development of QS-21 SMNP as a viable and safe vaccine adjuvant. Human use of QS-21 SMNP is currently being investigated with a germline targeting HIV vaccine N332-GT5 (ClinicalTrials.gov Identifier: NCT06033209).

Limitations: The study investigated different adjuvant doses, but not different immunogen doses. The HIV immunogen tested here is a model for difficult antigens with dense glycan shields. Long term durability of immune responses > 1-year post vaccination was not been studied here and needs to be investigated in future studies.

## Methods

### Sex as a biological variable

Our study examined male and female animals, and similar findings are reported for both sexes.

### Adjuvants

Saponins Quil-A (vac-quil) and QS-21 were purchased from InvivoGen and Desert King, respectively, and used as received. Phosphorylated HexaAcyl Disaccharide (PHAD®) MPLA was obtained from Avanti. Lab-scale SMNP were prepared as previously described (8). AS01_B_ was obtained from the Shingrix™ (GSK) vaccine vial that are separately provided from antigen. GMP-process QS-21 SMNP was prepared using a tangential flow filtration-based detergent dilution process (24). Briefly, SMNP components were solubilized in 20% (w/v) MEGA-10 surfactant at a 10:2:1:1 mass ratio of QS- 21:cholesterol:MPLA:dipalmitoylphosphatidyl choline at a batch size of 500 mg of QS-21. This starting detergent/lipids solution was diluted with phosphate buffered saline (PBS, pH 7.4) to reach a final QS- 21 concentration of 5 mg/mL and MEGA-10 concentration of 7.5%. The adjuvant/lipid mixture was then stepwise diluted with PBS (pH 7.4) with sufficient incubation time between each dilution step to allow for controlled SMNP self-assembly (**Supplemental Table 2**). After SMNP assembly, the sample was concentrated 20-fold on a 100 kDa hollow fiber TFF membrane (mPES, surface area: 0.16 m^2^, Repligen). After concentration, the sample underwent continuous diafiltration for 10 diafiltration volume equivalents against PBS pH 6.5. The sample was then filtered through a 0.2 µm membrane and used to fill 3 mL 2R Type I USP borosilicate glass vials with 0.6 mL of sample. Vials were closed with 13 mm bromobutyl 4023/50, Flurotec coated, B2-40 grey stoppers and a 13 mm aluminum caps with green plastic flip-off. Nanoparticle micrographs were acquired using Transmission Electron Microscopy (TEM) on a JEOL 2100F microscope (200 kV) with a magnification range of 10,000-60,000X. All images were recorded on a Gatan 2kx2k UltraScan CCD camera. Negative-stain sample preparation was performed by allowing 10 µL of SMNP to adsorb on a 200-mesh copper grid coated for one minute. The excess solution was then removed by touching the grid with a Kimwipe. The negative staining solution, 10 µL phosphotungstic acid (PTA) 1% aqueous, was then quickly added and the excess solution was removed with a Kimwipe. After drying at room temperature, the grid was mounted on a JEOL single tilt holder equipped in the TEM column for image capture.

### Protein production

BG505 MD39 untagged trimer immunogens and His-Avi-tagged biotinylated trimers [BG505 MD39 and BG505 MD39-base knockout (bKO)] (16) were produced by transient co-transfection of HEK-293F cells (Thermo Fisher Scientific) with Furin as previously described (25). His-Avi-tagged trimers were purified by immobilized metal ion affinity chromatography (IMAC) using HisTrap excel columns (Cytiva) followed by size-exclusion chromatography (SEC) using a Superdex 200 Increase 10/300 GL column (Cytiva). His-Avi-tagged trimers were biotinylated using BirA (Avidity) according to the manufacturer’s instructions and purified again to remove excess biotin using SEC. BG505 MD39 untagged trimer immunogens were purified by 2G12 affinity chromatography followed by SEC. Immunogen preps were confirmed to contain < 5 EU/mg of endotoxin using an Endosafe instrument (Charles River).

### Animals, immunizations and sample collections

Indian rhesus macaques (*Macaca mulatta*) were housed at the UC Davis California National Primate Research Center and maintained in accordance with NIH guidelines. Animal age, body weight and sex were equally distributed among groups. Animals ranged in age from 3-13 years (median of 5 years) and each group contained 3 females and 3 males (*n*=6 per group). Immunizations were prepared in sterile PBS with 100 µg BG505 MD39 HIV envelope immunogen and the appropriate amount of adjuvant.

Adjuvants administered differed in formulation, dose and delivery routes among groups. The following adjuvants with co-administration of 100 µg BG505 MD39 HIV immunogen were given: (i) Group 1 received 25 µg QS-21 SMNP, (ii) Group 2 received 50 µg QS-21 SMNP, (iii) Group 3 received 200 µg QS- 21 SMNP, (iv) Group 4 received 400 µg QS-21 SMNP, (v) Group 5 received 50 µg AS01_B_, (vi) Group 6 received 200 µg Quil-A SMNP, (vii) Group 7 received 200 µg QS-21 SMNP. The adjuvant dose relates to the quantity of saponin administered. The total amount of QS-21 is consistent between the 50 µg AS01_B_ and QS-21 SMNP formulations. Injections in groups 1-6 were given SC into the inner thigh (1 mL per injection) and Group 7 was given IM into the deltoid (0.5 mL per injection). Animals were immunized a total of 3 times at weeks 0, 10 and 24 as a unilateral bolus injection on the left side. All experimental manipulations (immunizations and sample collections) were performed under ketamine anesthesia (10 mg/kg body weight; IM). Blood samples were collected via peripheral venipuncture. Plasma and serum samples, peripheral blood mononuclear cells (PBMCs) and lymph node mononuclear cells (from FNAs) were processed and cryopreserved according to standard laboratory protocols, and subsequently shipped for downstream assays. Bone marrow aspirates were collected with heparinized syringes, placed in conical tubes containing R10 media and shipped overnight at 4°C for downstream assays.

### Detection of antigen-binding or total B_Mem_, B_GC_ and GC-T_FH_ by flow cytometry

LN FNAs or peripheral blood mononuclear cells (PBMCs) were thawed in RPMI media containing 10% FBS, 1x penicillin/streptomycin and 1x GlutaMAX. Biotinylated BG505 MD39 or BG505 MD39-base knockout (bKO) was tetramerized by fluorescently-conjugated streptavidin (SA). Biotinylated BG505 MD39 was tetramerized with SA-BV421 or SA-AF647. Biotinylated BG505 MD39-bKO was tetramerized with SA-PE. All SA-conjugates were added in a step-wise addition by adding 1/3 of the SA to the biotinylated protein at a time and incubating for 15 mins in between. PBMCs or LN FNAs were plated in a 96-well U-bottom plate at up to 5 x 10^6^ cells per well, stained with 1:25 Fc block (BD Biosciences, 564220) and washed with FACS buffer (2% FBS in PBS). Cells were first incubated with BG505 MD39- bKO probe for 20 minutes and then with BG505 MD39 probes for 30 minutes at 4°C in the dark. Cells were then stained with surface antibodies found in **Tables 1-3** depending on the sample type and incubated for an additional 30 minutes at 4°C in the dark. Following incubation, cells were washed twice with FACS buffer and acquired on a 5-laser equipped Aurora (Cytek Biosciences) spectral flow cytometer. Anti-CD38 PE-Cy5 was conjugated using purified anti-CD38 and the PE/Cy5 Conjugation Kit (Abcam, ab102893).

**Table 1.**
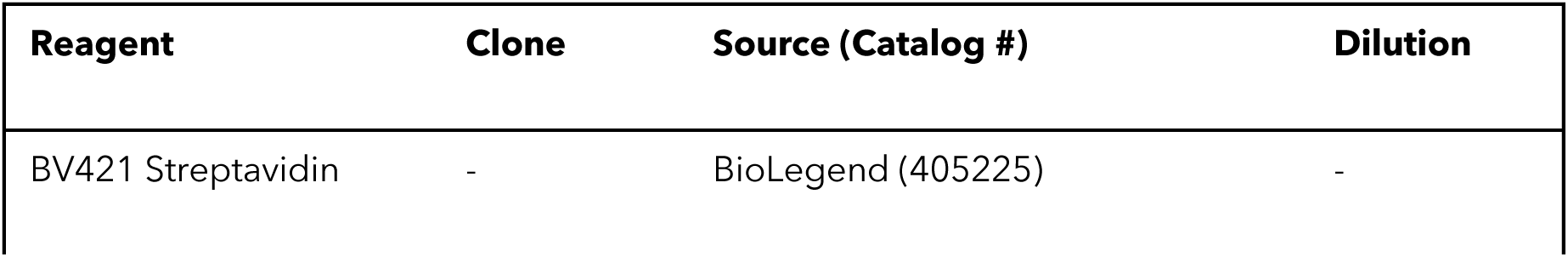
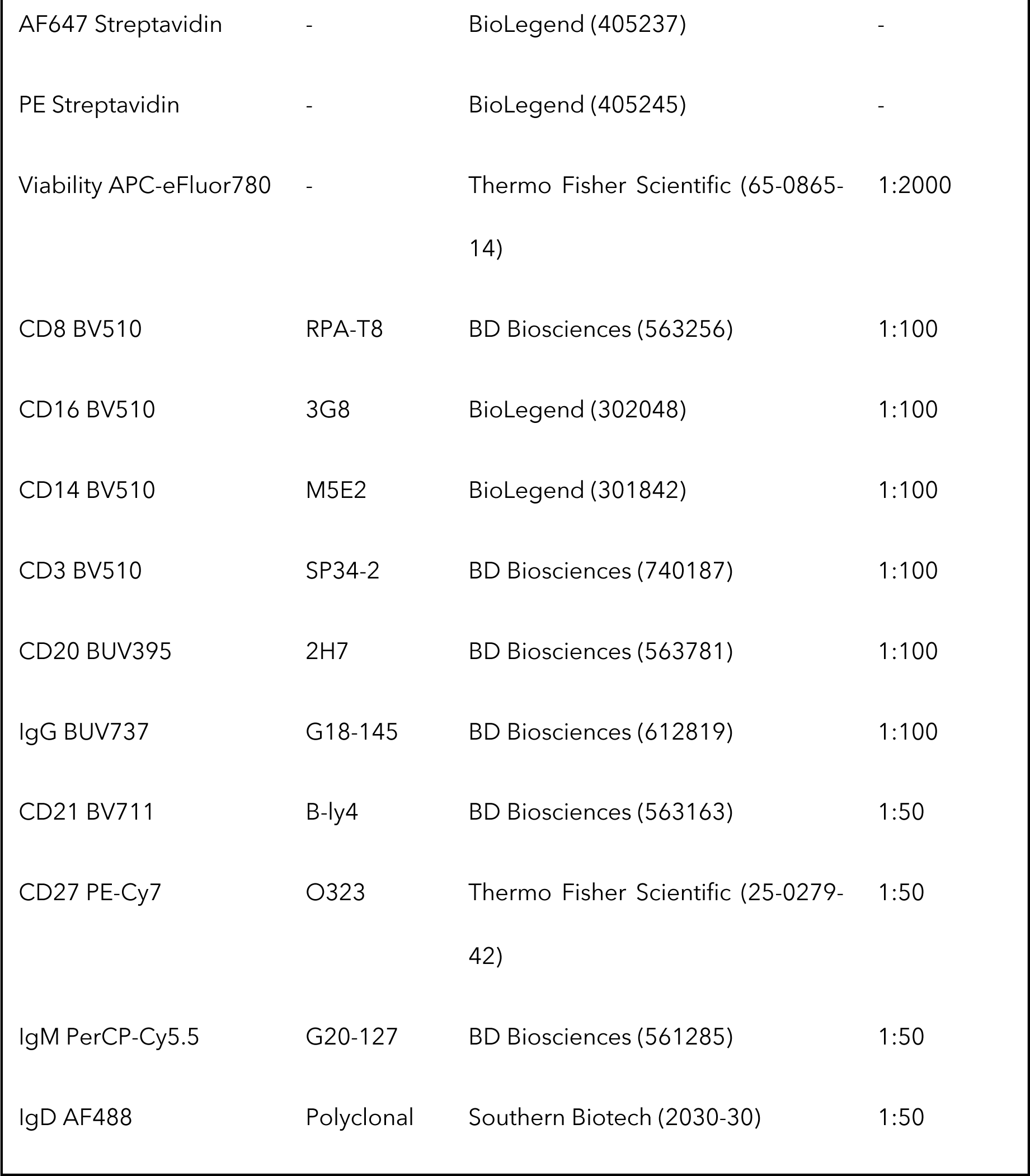
Staining panel for antigen-specific B_Mem_ in PBMCs.

| Reagent | Clone | Source (Catalog #) | Dilution |
| --- | --- | --- | --- |
| BV421 Streptavidin | - | BioLegend (405225) | - |
| AF647 Streptavidin | - | BioLegend (405237) | - |
| PE Streptavidin | - | BioLegend (405245) | - |
| Viability APC-eFluor780 | - | Thermo Fisher Scientific (65-0865-14) | 1:2000 |
| CD8 BV510 | RPA-T8 | BD Biosciences (563256) | 1:100 |
| CD16 BV510 | 3G8 | BioLegend (302048) | 1:100 |
| CD14 BV510 | M5E2 | BioLegend (301842) | 1:100 |
| CD3 BV510 | SP34-2 | BD Biosciences (740187) | 1:100 |
| CD20 BUV395 | 2H7 | BD Biosciences (563781) | 1:100 |
| IgG BUV737 | G18-145 | BD Biosciences (612819) | 1:100 |
| CD21 BV711 | B-ly4 | BD Biosciences (563163) | 1:50 |
| CD27 PE-Cy7 | O323 | Thermo Fisher Scientific (25-0279-42) | 1:50 |
| IgM PerCP-Cy5.5 | G20-127 | BD Biosciences (561285) | 1:50 |
| IgD AF488 | Polyclonal | Southern Biotech (2030-30) | 1:50 |

**Table 2.**
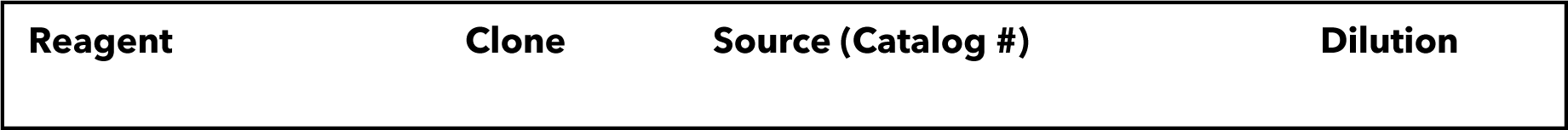
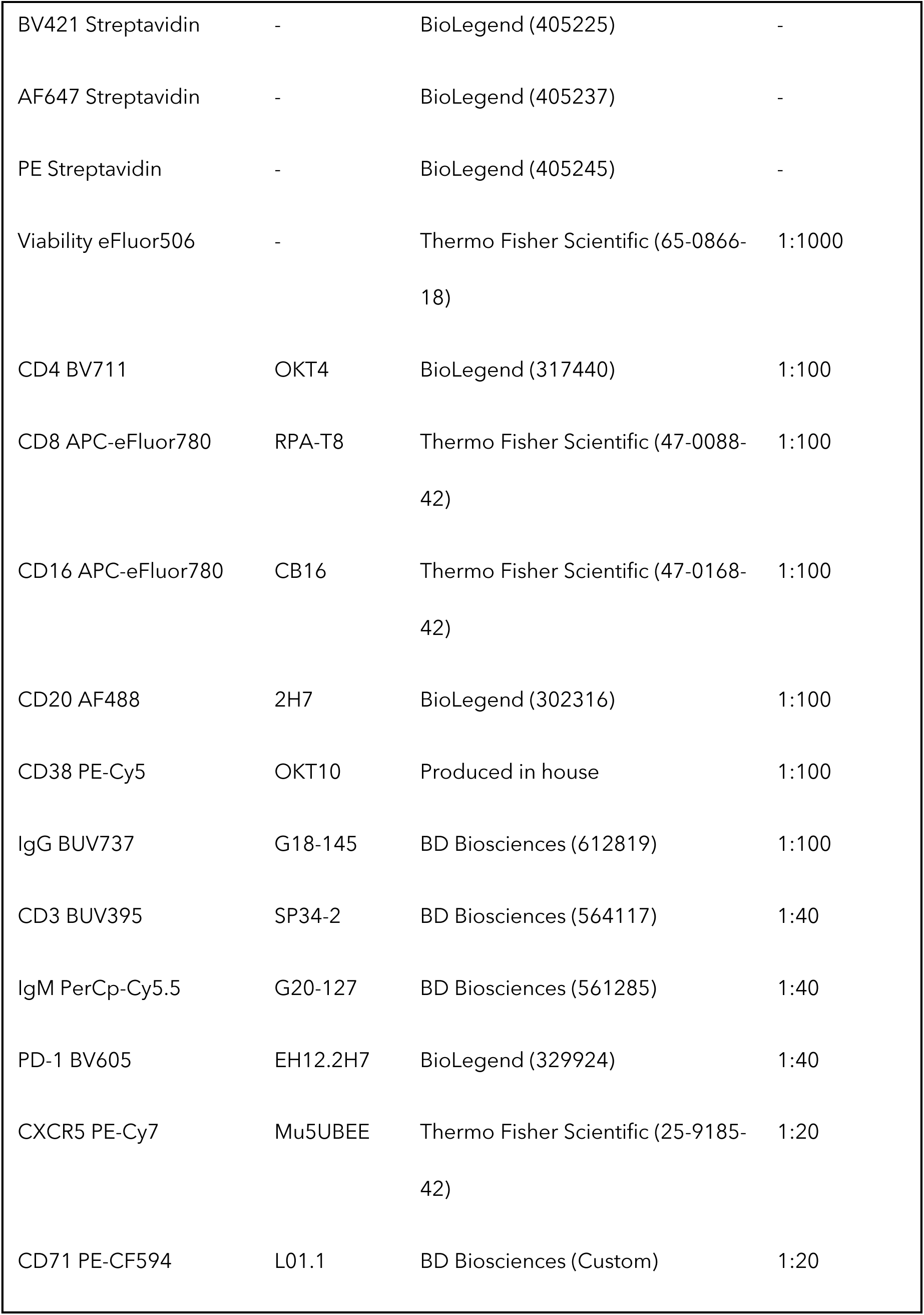
Staining panel for antigen-specific B_GC_ and B_Mem_ in ipsilateral LN FNAs.

| Reagent | Clone | Source (Catalog #) | Dilution |
| --- | --- | --- | --- |
| BV421 Streptavidin | - | BioLegend (405225) | - |
| AF647 Streptavidin | - | BioLegend (405237) | - |
| PE Streptavidin | - | BioLegend (405245) | - |
| Viability eFluor506 | - | Thermo Fisher Scientific (65-0866-18) | 1:1000 |
| CD4 BV711 | OKT4 | BioLegend (317440) | 1:100 |
| CD8 APC-eFluor780 | RPA-T8 | Thermo Fisher Scientific (47-0088-42) | 1:100 |
| CD16 APC-eFluor780 | CB16 | Thermo Fisher Scientific (47-0168-42) | 1:100 |
| CD20 AF488 | 2H7 | BioLegend (302316) | 1:100 |
| CD38 PE-Cy5 | OKT10 | Produced in house | 1:100 |
| IgG BUV737 | G18-145 | BD Biosciences (612819) | 1:100 |
| CD3 BUV395 | SP34-2 | BD Biosciences (564117) | 1:40 |
| IgM PerCp-Cy5.5 | G20-127 | BD Biosciences (561285) | 1:40 |
| PD-1 BV605 | EH12.2H7 | BioLegend (329924) | 1:40 |
| CXCR5 PE-Cy7 | Mu5UBEE | Thermo Fisher Scientific (25-9185-42) | 1:20 |
| CD71 PE-CF594 | L01.1 | BD Biosciences (Custom) | 1:20 |

**Table 3.** Staining panel for antigen-specific B_GC_ and B_Mem_ in contralateral LN FNAs.

| Reagent | Clone | Source (Catalog #) | Dilution |
| --- | --- | --- | --- |
| BV421 Streptavidin | - | BioLegend (405225) | - |
| AF647 Streptavidin | - | BioLegend (405237) | - |
| PE Streptavidin | - | BioLegend (405245) | - |
| Viability APC-eFluor780 | - | Thermo Fisher Scientific (65-0865-14) | 1:2000 |
| CD8 BV510 | RPA-T8 | BD Biosciences (563256) | 1:100 |
| CD16 BV510 | 3G8 | BioLegend (302048) | 1:100 |
| CD14 BV510 | M5E2 | BioLegend (301842) | 1:100 |
| CD3 BV510 | SP34-2 | BD Biosciences (569486) | 1:100 |
| CD20 BUV395 | 2H7 | BD Biosciences (563782) | 1:100 |
| IgG BUV737 | G18-145 | BD Biosciences (612819) | 1:100 |
| CD38 PE-Cy5 | OKT10 | Produced in house | 1:200 |
| CD27 PE-Cy7 | O323 | Thermo Fisher Scientific (25-0279-42) | 1:50 |
| CD21 BV711 | B-ly4 | BD Biosciences (563163) | 1:50 |
| IgM PerCP-Cy5.5 | G20-127 | BD Biosciences (561285) | 1:50 |
| IgD AF488 | Polyclonal | Southern Biotech (2030-30) | 1:50 |
| CD69 BV650 | FN50 | BD Biosciences (563835) | 1:100 |
| CD71 PE-CF594 | L01.1 | BD Biosciences (Custom) | 1:20 |

For analysis of bulk B_GC_ and GC-T_FH_ a threshold of 250 B cells and 500 CD4^+^ T cells was used, respectively. For analysis of Env-binding B_GC_, a threshold of 75 B_GC_ cells was used. For Env-binding B_Mem_ in LN FNAs, samples with at least 1000 B cells were included for analysis. For analysis of bKO^+^ Env- binding B_Mem_ in the blood, a threshold of 10 Env^+^ B_Mem_ was used. The limit of detection (LOD) for Env- binding B_GC_ and B_Mem_ in LN FNAs was 0.01% and was determined by calculating the median of (3/(number of B cells recorded)) from pre-immunized samples. The LOD for Env-binding B_Mem_ in the blood was 0.0006% and was determined by calculating the median + 2x standard deviation of (1/(number of B cells recorded)) from pre-immunized samples. Any sample that met the above threshold criteria and fell below 0.001% was set to a baseline of 0.001%. To calculate B_Mem_ half-life (*t*_1/2_) between week 12 and 16 or week 16 and 24, data was log_2_ transformed and a simple linear regression was utilized.

### Single B cell sequencing

Env-binding B_Mem_ at week 12 from 50 µg and 400 µg QS-21 SMNP groups were stained with tetramerized BG505 MD39 probes using TotalSeq-C oligo-tagged fluorescent-SA, surface antibodies and TotalSeq-C oligo-tagged anti-β2M & CD298 antibodies to multiplex animal samples (**Table 4**). Samples were sorted on a FACSymphony S6 (BD Biosciences) into 1.5 mL tubes with FBS supplemented RPMI media.

**Table 4.** Staining panel for sorting antigen-specific B_Mem_ in PBMCs.

| Reagent | Clone | Source (Catalog #) | Dilution |
| --- | --- | --- | --- |
| BV421 Streptavidin | - | BioLegend (custom) | - |
| AF647 Streptavidin | - | BioLegend (405237) | - |
| PE Streptavidin | - | BioLegend (405261) | - |
| Viability APC-eFluor780 | - | Thermo Fisher Scientific (65-0865-14) | 1:2000 |
| CD8 BV510 | RPA-T8 | BD Biosciences (563256) | 1:100 |
| CD16 BV510 | 3G8 | BioLegend (302048) | 1:100 |
| CD14 BV510 | M5E2 | BioLegend (301842) | 1:100 |
| CD3 BV510 | SP34-2 | BD Biosciences (569486) | 1:100 |
| CD20 BUV395 | 2H7 | BD Biosciences (563782) | 1:100 |
| IgG BV605 | G18-145 | BD Biosciences (563246) | 1:100 |
| IgM PerCP-Cy5.5 | G20-127 | BD Biosciences (561285) | 1:50 |
| IgD AF488 | Polyclonal | Southern Biotech (2030-30) | 1:50 |
| CD27 PE-Cy7 | O323 | Thermo Fisher Scientific (25-0279-42) | 1:50 |

After sorting, samples were washed once with PBS and loaded onto a Chromium chip and controller according to manufacturer’s recommendation (10X Genomics). VDJ, GEX and HTO libraries were generated and sequenced on a NovaSeq 6000 (Illumina). Cell Ranger v7.2.0 (10X Genomics) was used to demultiplex sequencing files into FASTQ files and assemble reads from the libraries. VDJ reads were assembled de novo in Cell Ranger and contigs were aligned to a custom *Macaca mulatta* germline VDJ reference using IgBLAST and the Change-O 1.3.0 package from the Immcantation framework (46). Sequences were assigned to animals using the MULTIseqDemux function in Seurat v5 (47). Inferred germline sequences (CreateGermline.py) and clones (DefineClones.py) were determined with Change-O. SHM were quantified using the observedMutations command in SHazaM package v1.2.0 comparing HC or LC sequences to the inferred germline sequences (masking the D gene sequences). For clonal abundance, clonotypes for each animal were quantified using the countClones function in Alakazam package v1.3.0. Clonal richness (Chao1) was calculated using the iNEXT package v3.0.0 in R (48). Gene expression integrative analysis was done with Seurat v5 using CCAIntegration. Cells expressing less than 200 or more than 4500 transcripts and more than 5% mitochondrial genes were excluded. Clusters containing non-B cells were removed for analysis.

### Detection of antigen-specific CD4 or CD8 T cells by flow cytometry

Activation-induced marker (AIM) and intracellular cytokine staining (ICS) assay for the detection of antigen-specific T cells was performed as previously described (20). Cryopreserved PBMCs were thawed and washed in RPMI media containing 10% FBS, 1x penicillin/streptomycin and 1x GlutaMAX (R10 media). Cells were counted and plated at 1 × 10^6^ cells per well in a round-bottom 96-well plate. Prior to stimulation, cells were blocked with 0.5 μg/mL anti-CD40 mAb (Miltenyi Biotec) and stained with anti- CXCR5 and CCR7 for 15 minutes at 37°C. Then, cells were incubated for 24 hours in the presence of: (1) 5 µg/mL BG505 MD39 Env peptide pool; (2) 1 ng/mL staphylococcal enterotoxin B (SEB) used as a positive control; or (3) an equimolar amount of DMSO as negative, unstimulated control plated in duplicate. BG505 MD39 Env peptide pools consist of overlapping 15-mer peptides that cover the entire protein sequence and were resuspended in DMSO. After 24 hours of incubation, intracellular transport inhibitors – 0.25 μL/well of GolgiPlug (BD Biosciences) and 0.25 μL/well of GolgiStop (BD Biosciences) – and AIM marker antibodies (CD25, CD40L, CD69, OX40, 4-1BB) were added to the samples and incubated for another 4 hours. Cells were then centrifuged and incubated for 15 min at RT with Fc Block (BioLegend) and Fixable Live/Dead Blue (Invitrogen). Cells were then washed with FACS buffer (2% FBS in PBS) and stained with the surface antibodies for 30 minutes at 4°C. Following, cells were fixed with 4% formaldehyde, permeabilized with a saponin-based buffer and stained with intracellular cytokine antibodies for 30 minutes at RT. Cells were washed with PBS and stored at 4°C until acquired a 5-laser equipped Aurora (Cytek Biosciences) spectral flow cytometer. Antibody panels and reagents are summarized in **Table 5**. For data analysis, antigen-specific T cells were measured as background subtracted data, where the linear averages of the DMSO background signal, calculated from duplicate wells for each sample, were deducted from the stimulated signal. A minimum threshold for DMSO background signals was set at 0.005% and the limit of quantitation (LOQ) was defined as the geometric mean of all DMSO negative control wells. For each sample, the stimulation index (SI) was calculated as the ratio between the AIM^+^ response in the stimulated condition and the average DMSO response for the same sample. Samples with an SI lower than 2 for CD4 or 3 for CD8 T cells responses and/or with a background subtracted response lower than the LOQ were considered as non-responders. Non-responder samples were set to 2-fold below the LOQ. Sample 48005 from Group 1 at week 2, was excluded from antigen-specific T cell analysis due to the viability of the cells.

**Table 5.**
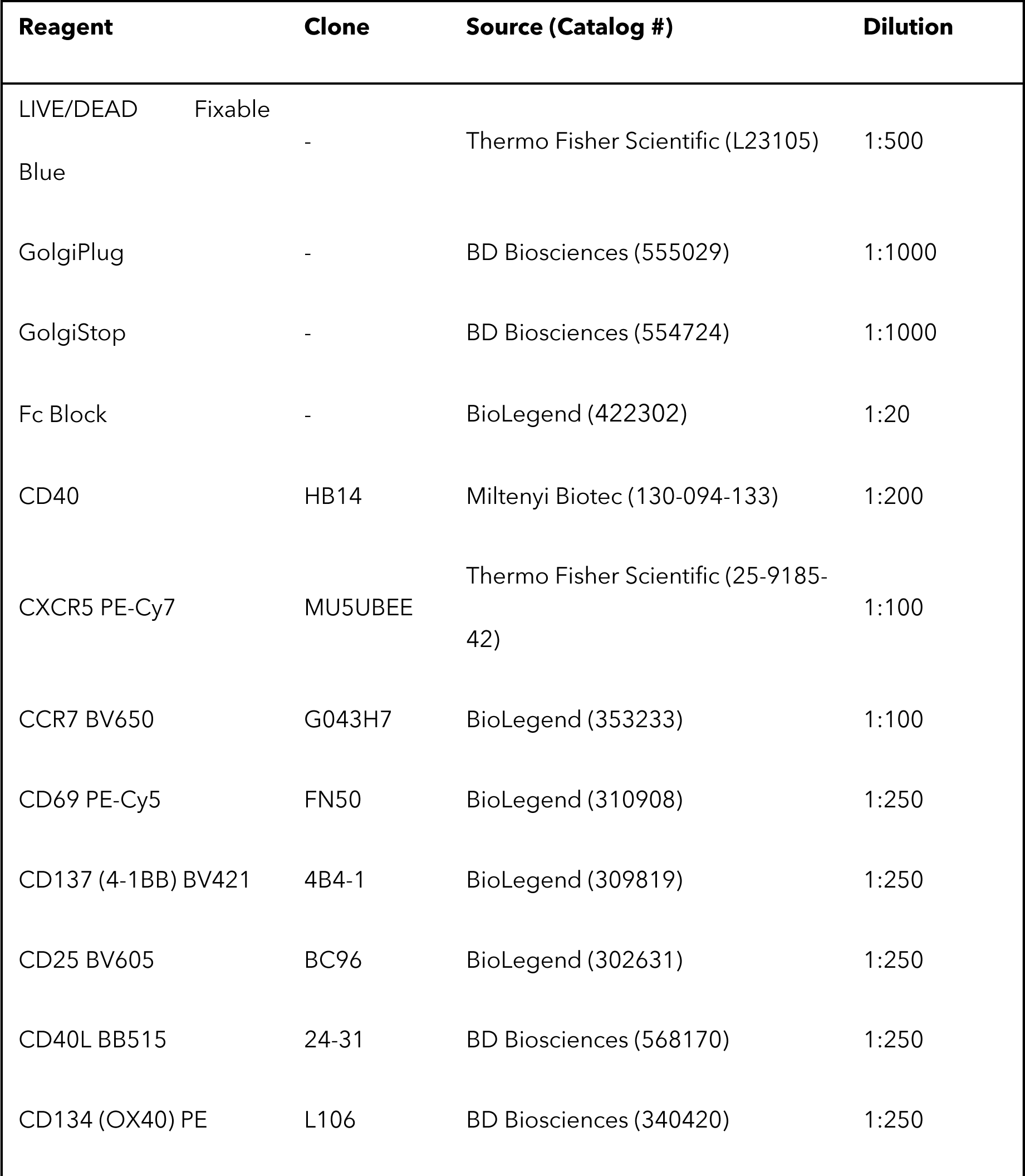
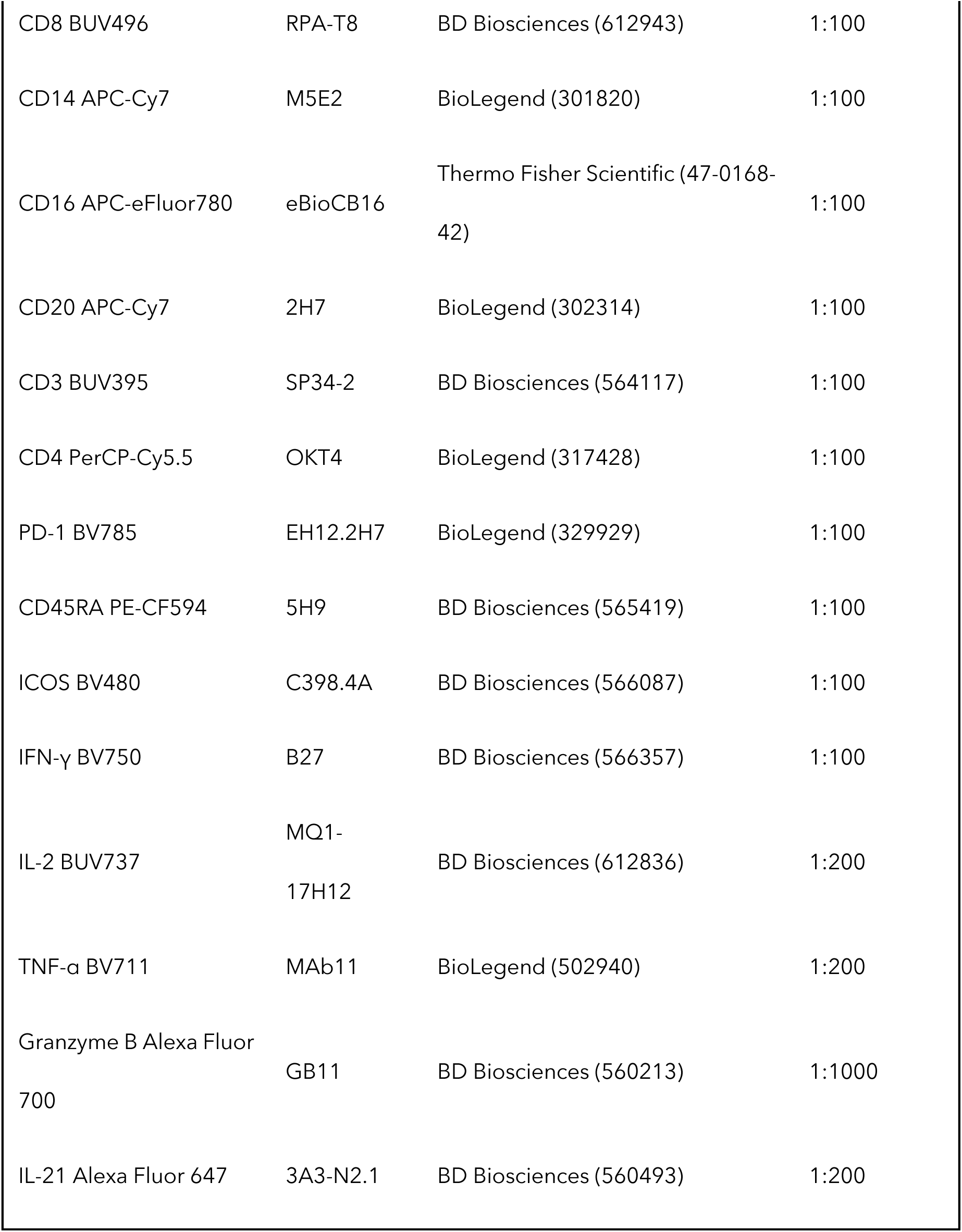
AIM/ICS staining panel for antigen-specific T cells in PBMCs.

### ELISA

Plasma samples from animals were serially diluted (starting at 1:50) and plated on pre-coated wells with BG505 MD39 antigen. Fort this, 96-well plates (Corning Cat# 3690) were coated with SA (Thermo Fisher Scientific Cat# 434302) first, blocked with 2% BSA and then incubated with biotinylated BG505 MD39. Env binding-IgG antibodies were detected using HRP goat anti-human IgG (Jackson ImmunoResearch Cat# 109-035-098) and 1-Step™ Ultra TMB ELISA Substrate Solution (Thermo Fisher Scientific Cat# 34029). The reaction was stopped with 2N of sulfuric acid (Ricca Chemical Cat# 8310-32) and read at 450 nm on a FlexStation 3 plate reader (Molecular Devices).

### Neutralization assays

Neutralization assays using the BG505.W6M.ENV.C2 isolate (T332N) pseudovirus were performed as previously described (42). Assays were run blinded and in duplicates. Initial serum dilution was set at 1:10 and values below were considered as non-neutralizing. Absolute ID_50_ values were calculated using normalized relative luminescence units and a customized nonlinear regression model with the bottom constraint set to 0 and top constraint set to the <100:

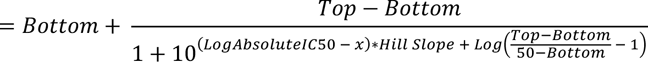

### ELISPOT

Bone marrow aspirates were processed to harvest mononuclear cells which were serially diluted and plated onto pre-coated wells with BG505 MD39 antigen (bound via Galanthus Nivalis Lectin [Vector laboratories Cat# L-1240]) or total goat anti-human Ig antibody (Southern Biotech Cat# 2010-01) as captures. Plates were incubated in a 5% CO_2_ incubator at 37°C for 16 hours. Secreted antibodies were detected using biotinylated goat anti-human IgG (Southern Biotech Cat# 2045-08), HRP Avidin D (Vector laboratories Cat# A-2004) and AEC substrate (Acros Organics Cat# 147870250). Formed spots were imaged using the Immunospot CTL counter and Image Acquisition 4.5 software (Cellular Technology), and counted manually.

### Statistics

Statistical tests were calculated in GraphPad Prism 10. An unpaired two-tailed Mann-Whitney test was used for the following comparisons: (i) Group 1 vs 2, Group 2 vs 3, Group 3 vs 4, Group 1 vs 4, (ii) Group 5 vs Group 2, (iii) Group 6 vs Group 3 and (iv) Group 7 vs Group 3. A Spearman’s Correlation test was used to analyze all correlations. *P* values are defined as follows: not significant (NS) *P* > 0.05, \**P* ≤ 0.05, \*\**P* < 0.01, \*\*\**P* < 0.001, \*\*\*\**P* < 0.0001.

### Study approval

All procedures of animal husbandry and sample collections were performed according to established Standard Operating Procedures that were approved by the Institutional Animal Care and Use Committee of the University of California, Davis, and that were in compliance with the 2011 “*Guide for the Care and Use of Laboratory Animals”*.

## Data availability

All data required for the conclusions of this work are included in the main figures or the supplemental information.

## Author contributions

DJI and SC designed the original animal study. KKAVR and PRR adapted and supervised the animal study. DJI, SC, and PRR designed the majority of the experiments. PRR, AM, and NIB performed and analyzed B cell responses in the blood and LNs. EMZ designed, performed and analyzed T cell assays with technical assistance from PGL. LM and KKM performed and analyzed plasma cytokine and antibody responses. ISP performed and analyzed TEM. IB, LH, and DRB performed and analyzed serum neutralization assays. AP and SPK performed and analyzed bone marrow plasma cell responses. WRS, CAC and BG supplied immunogens and reagents. SP, ES, and SLS supervised GMP-process SMNP production. AW, GS, MB, and DK supplied GMP-process SMNP. PRR wrote the original draft with substantial input from SC, DJI and AM. WRS, DRB, EMZ, KKAVR, SPK, AP, and CAC edited the manuscript. All authors read and approved the manuscript.

## Acknowledgments

We thank Jennifer Watanabe, Jodie Usachenko, and the staff of Primate Services and Clinical Laboratories of the California National Primate Research Center for technical assistance. We thank Diane G Carnathan, Sherrie Jean and Jennifer S Wood for guidance and feedback on LN FNAs. We are grateful to the Flow Cytometry and Next Generation Sequencing core facilities at La Jolla Institute for Immunology for their services. This work was funded by the National Institute of Allergy and Infectious Diseases of the NIH (NIAID-NIH) under award CHAVD UM1AI144462 and from the Office of Research Infrastructure Program, Office of the Director, NIH to the California National Primate Research Center under award 51OD011107. The NovaSeq 6000 used in this work was funded by NIH grant S10OD025052.

## References

1. Del Giudice G, Rappuoli R, and Didierlaurent AM. Correlates of adjuvanticity: A review on adjuvants in licensed vaccines. Semin Immunol. 2018;39:14–21.

2. Pulendran B, P SA, and O’Hagan DT. Emerging concepts in the science of vaccine adjuvants. Nat Rev Drug Discov. 2021;20(6):454–75.

3. Singh A, Boggiano C, Eller MA, Maciel M, Jr., Marovich MA, Mehra VL, et al. Optimizing the Immunogenicity of HIV Vaccines by Adjuvants - NIAID Workshop Report. Vaccine. 2023;41(31):4439–46.

4. Crotty S. T Follicular Helper Cell Biology: A Decade of Discovery and Diseases. Immunity. 2019;50(5):1132–48.

5. Garçon N, Morel S, Didierlaurent A, Descamps D, Wettendorff M, and Van Mechelen M. Development of an AS04-adjuvanted HPV vaccine with the adjuvant system approach. BioDrugs. 2011;25(4):217–26.

6. Heyward WL, Kyle M, Blumenau J, Davis M, Reisinger K, Kabongo ML, et al. Immunogenicity and safety of an investigational hepatitis B vaccine with a Toll-like receptor 9 agonist adjuvant (HBsAg-1018) compared to a licensed hepatitis B vaccine in healthy adults 40-70 years of age. Vaccine. 2013;31(46):5300–5.

7. Kasturi SP, Rasheed MAU, Havenar-Daughton C, Pham M, Legere T, Sher ZJ, et al. 3M-052, a synthetic TLR-7/8 agonist, induces durable HIV-1 envelope-specific plasma cells and humoral immunity in nonhuman primates. Sci Immunol. 2020;5(48).

8. Silva M, Kato Y, Melo MB, Phung I, Freeman BL, Li Z, et al. A particulate saponin/TLR agonist vaccine adjuvant alters lymph flow and modulates adaptive immunity. Sci Immunol. 2021;6(66):eabf1152.

9. Burny W, Callegaro A, Bechtold V, Clement F, Delhaye S, Fissette L, et al. Different Adjuvants Induce Common Innate Pathways That Are Associated with Enhanced Adaptive Responses against a Model Antigen in Humans. Front Immunol. 2017;8:943.

10. Leroux-Roels G, Marchant A, Levy J, Van Damme P, Schwarz TF, Horsmans Y, et al. Impact of adjuvants on CD4(+) T cell and B cell responses to a protein antigen vaccine: Results from a phase II, randomized, multicenter trial. Clin Immunol. 2016;169:16–27.

11. Agnandji ST, Lell B, Fernandes JF, Abossolo BP, Methogo BG, Kabwende AL, et al. A phase 3 trial of RTS,S/AS01 malaria vaccine in African infants. N Engl J Med. 2012;367(24):2284–95.

12. Lal H, Cunningham AL, Godeaux O, Chlibek R, Diez-Domingo J, Hwang SJ, et al. Efficacy of an adjuvanted herpes zoster subunit vaccine in older adults. N Engl J Med. 2015;372(22):2087–96.

13. Lenart K, Arcoverde Cerveira R, Hellgren F, Ols S, Sheward DJ, Kim C, et al. Three immunizations with Novavax’s protein vaccines increase antibody breadth and provide durable protection from SARS-CoV-2. NPJ Vaccines. 2024;9(1):17.

14. Lee JH, and Crotty S. HIV vaccinology: 2021 update. Semin Immunol. 2021;51:101470.

15. Haynes BF, Wiehe K, Borrow P, Saunders KO, Korber B, Wagh K, et al. Strategies for HIV-1 vaccines that induce broadly neutralizing antibodies. Nat Rev Immunol. 2023;23(3):142–58.

16. Lee JH, Sutton HJ, Cottrell CA, Phung I, Ozorowski G, Sewall LM, et al. Long-primed germinal centres with enduring affinity maturation and clonal migration. Nature. 2022;609(7929):998–1004.

17. Steichen JM, Lin YC, Havenar-Daughton C, Pecetta S, Ozorowski G, Willis JR, et al. A generalized HIV vaccine design strategy for priming of broadly neutralizing antibody responses. Science. 2019;366(6470).

18. Steichen JM, Phung I, Salcedo E, Ozorowski G, Willis JR, Baboo S, et al. Vaccine priming of rare HIV broadly neutralizing antibody precursors in nonhuman primates. Science. 2024;384(6697):eadj8321.

19. Schiffner T, Phung I, Ray R, Irimia A, Tian M, Swanson O, et al. Vaccination induces broadly neutralizing antibody precursors to HIV gp41. Nat Immunol. 2024;25(6):1073–82.

20. Phung I, Rodrigues KA, Marina-Zárate E, Maiorino L, Pahar B, Lee WH, et al. A combined adjuvant approach primes robust germinal center responses and humoral immunity in non-human primates. Nat Commun. 2023;14(1):7107.

21. Tarantal AF, Noctor SC, and Hartigan-O’Connor DJ. Nonhuman Primates in Translational Research. Annu Rev Anim Biosci. 2022;10:441–68.

22. Sun HX, Xie Y, and Ye YP. ISCOMs and ISCOMATRIX. Vaccine. 2009;27(33):4388–401.

23. Reimer JM, Karlsson KH, Lövgren-Bengtsson K, Magnusson SE, Fuentes A, and Stertman L. Matrix-M™ adjuvant induces local recruitment, activation and maturation of central immune cells in absence of antigen. PLoS One. 2012;7(7):e41451.

24. Pires IS, Ni K, Melo MB, Li N, Ben-Akiva E, Maiorino L, et al. Controlled lipid self-assembly for scalable manufacturing of next-generation immune stimulating complexes. Chemical Engineering Journal. 2023;464:142664.

25. Steichen JM, Kulp DW, Tokatlian T, Escolano A, Dosenovic P, Stanfield RL, et al. HIV Vaccine Design to Target Germline Precursors of Glycan-Dependent Broadly Neutralizing Antibodies. Immunity. 2016;45(3):483–96.

26. Abbott RK, and Crotty S. Factors in B cell competition and immunodominance. Immunol Rev. 2020;296(1):120–31.

27. Cirelli KM, Carnathan DG, Nogal B, Martin JT, Rodriguez OL, Upadhyay AA, et al. Slow Delivery Immunization Enhances HIV Neutralizing Antibody and Germinal Center Responses via Modulation of Immunodominance. Cell. 2019;177(5):1153–71.e28.

28. Havenar-Daughton C, Carnathan DG, Torrents de la Peña A, Pauthner M, Briney B, Reiss SM, et al. Direct Probing of Germinal Center Responses Reveals Immunological Features and Bottlenecks for Neutralizing Antibody Responses to HIV Env Trimer. Cell Rep. 2016;17(9):2195–209.

29. Locci M, Havenar-Daughton C, Landais E, Wu J, Kroenke MA, Arlehamn CL, et al. Human circulating PD-1+CXCR3-CXCR5+ memory Tfh cells are highly functional and correlate with broadly neutralizing HIV antibody responses. Immunity. 2013;39(4):758–69.

30. Morita R, Schmitt N, Bentebibel SE, Ranganathan R, Bourdery L, Zurawski G, et al. Human blood CXCR5(+)CD4(+) T cells are counterparts of T follicular cells and contain specific subsets that differentially support antibody secretion. Immunity. 2011;34(1):108–21.

31. Heit A, Schmitz F, Gerdts S, Flach B, Moore MS, Perkins JA, et al. Vaccination establishes clonal relatives of germinal center T cells in the blood of humans. J Exp Med. 2017;214(7):2139–52.

32. Vella LA, Buggert M, Manne S, Herati RS, Sayin I, Kuri-Cervantes L, et al. T follicular helper cells in human efferent lymph retain lymphoid characteristics. J Clin Invest. 2019;129(8):3185–200.

33. Nielsen CM, Ogbe A, Pedroza-Pacheco I, Doeleman SE, Chen Y, Silk SE, et al. Protein/AS01(B) vaccination elicits stronger, more Th2-skewed antigen-specific human T follicular helper cell responses than heterologous viral vectors. Cell Rep Med. 2021;2(3):100207.

34. Zhang Z, Mateus J, Coelho CH, Dan JM, Moderbacher CR, Gálvez RI, et al. Humoral and cellular immune memory to four COVID-19 vaccines. Cell. 2022;185(14):2434–51.e17.

35. Rydyznski Moderbacher C, Kim C, Mateus J, Plested J, Zhu M, Cloney-Clark S, et al. NVX-CoV2373 vaccination induces functional SARS-CoV-2-specific CD4+ and CD8+ T cell responses. J Clin Invest. 2022;132(19).

36. U.S. Food and Drug Administration; 2023.

37. Feng Y, Yuan M, Powers JM, Hu M, Munt JE, Arunachalam PS, et al. Broadly neutralizing antibodies against sarbecoviruses generated by immunization of macaques with an AS03-adjuvanted COVID-19 vaccine. Sci Transl Med. 2023;15(695):eadg7404.

38. Muecksch F, Wang Z, Cho A, Gaebler C, Ben Tanfous T, DaSilva J, et al. Increased memory B cell potency and breadth after a SARS-CoV-2 mRNA boost. Nature. 2022;607(7917):128–34.

39. Leggat DJ, Cohen KW, Willis JR, Fulp WJ, deCamp AC, Kalyuzhniy O, et al. Vaccination induces HIV broadly neutralizing antibody precursors in humans. Science. 2022;378(6623):eadd6502.

40. Turner JS, Zhou JQ, Han J, Schmitz AJ, Rizk AA, Alsoussi WB, et al. Human germinal centres engage memory and naive B cells after influenza vaccination. Nature. 2020;586(7827):127–32.

41. Didierlaurent AM, Laupèze B, Di Pasquale A, Hergli N, Collignon C, and Garçon N. Adjuvant system AS01: helping to overcome the challenges of modern vaccines. Expert Rev Vaccines. 2017;16(1):55–63.

42. Pauthner M, Havenar-Daughton C, Sok D, Nkolola JP, Bastidas R, Boopathy AV, et al. Elicitation of Robust Tier 2 Neutralizing Antibody Responses in Nonhuman Primates by HIV Envelope Trimer Immunization Using Optimized Approaches. Immunity. 2017;46(6):1073–88.e6.

43. Ols S, Yang L, Thompson EA, Pushparaj P, Tran K, Liang F, et al. Route of Vaccine Administration Alters Antigen Trafficking but Not Innate or Adaptive Immunity. Cell Rep. 2020;30(12):3964–71.e7.

44. Martin LBB, Kikuchi S, Rejzek M, Owen C, Reed J, Orme A, et al. Complete biosynthesis of the potent vaccine adjuvant QS-21. Nat Chem Biol. 2024;20(4):493–502.

45. Liu Y, Zhao X, Gan F, Chen X, Deng K, Crowe SA, et al. Complete biosynthesis of QS-21 in engineered yeast. Nature. 2024;629(8013):937–44.

46. Gupta NT, Vander Heiden JA, Uduman M, Gadala-Maria D, Yaari G, and Kleinstein SH. Change-O: a toolkit for analyzing large-scale B cell immunoglobulin repertoire sequencing data. Bioinformatics. 2015;31(20):3356–8.

47. Hao Y, Stuart T, Kowalski MH, Choudhary S, Hoffman P, Hartman A, et al. Dictionary learning for integrative, multimodal and scalable single-cell analysis. Nat Biotechnol. 2024;42(2):293–304.

48. Hsieh TC, Ma KH, and Chao A. iNEXT: an R package for rarefaction and extrapolation of species diversity (Hill numbers). Methods in Ecology and Evolution. 2016;7(12):1451–6.

