## Supplementary Material for "Dose-dependent regulation of immune memory responses against HIV by saponin monophosphoryl lipid A nanoparticle adjuvant"

**FigS1**

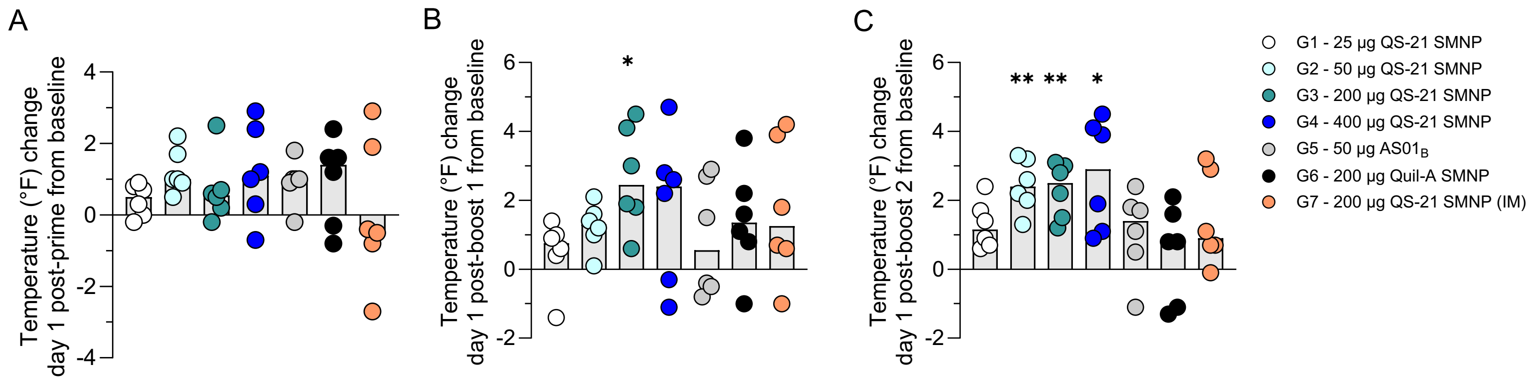
A
B
C
Temperature (°F) change day 1 post-prime from baseline
Temperature (°F) change day 1 post-boost 1 from baseline
Temperature (°F) change day 1 post-boost 2 from baseline
*
** ** *
G1 - 25 µg QS-21 SMNP
G2 - 50 µg QS-21 SMNP
G3 - 200 µg QS-21 SMNP
G4 - 400 µg QS-21 SMNP
G5 - 50 µg AS01B
G6 - 200 µg Quil-A SMNP
G7 - 200 µg QS-21 SMNP (IM)

**Supplemental Figure 1. Animal body temperature changes post-immunizations.**
**(A-C)** Change in rectal body temperature (°F) from baseline to day 1 **(A)** post-prime, **(B)** post-boost 1 and **(C)** post-boost 2.
Gray bars represent the median. Data was analyzed using a two-way ANOVA comparing baseline temperatures to temperature post-injection within each group. $^{*}P \leq 0.05$, $^{**}P < 0.01$, $^{***}P < 0.001$, $^{****}P < 0.0001$.

FigS2

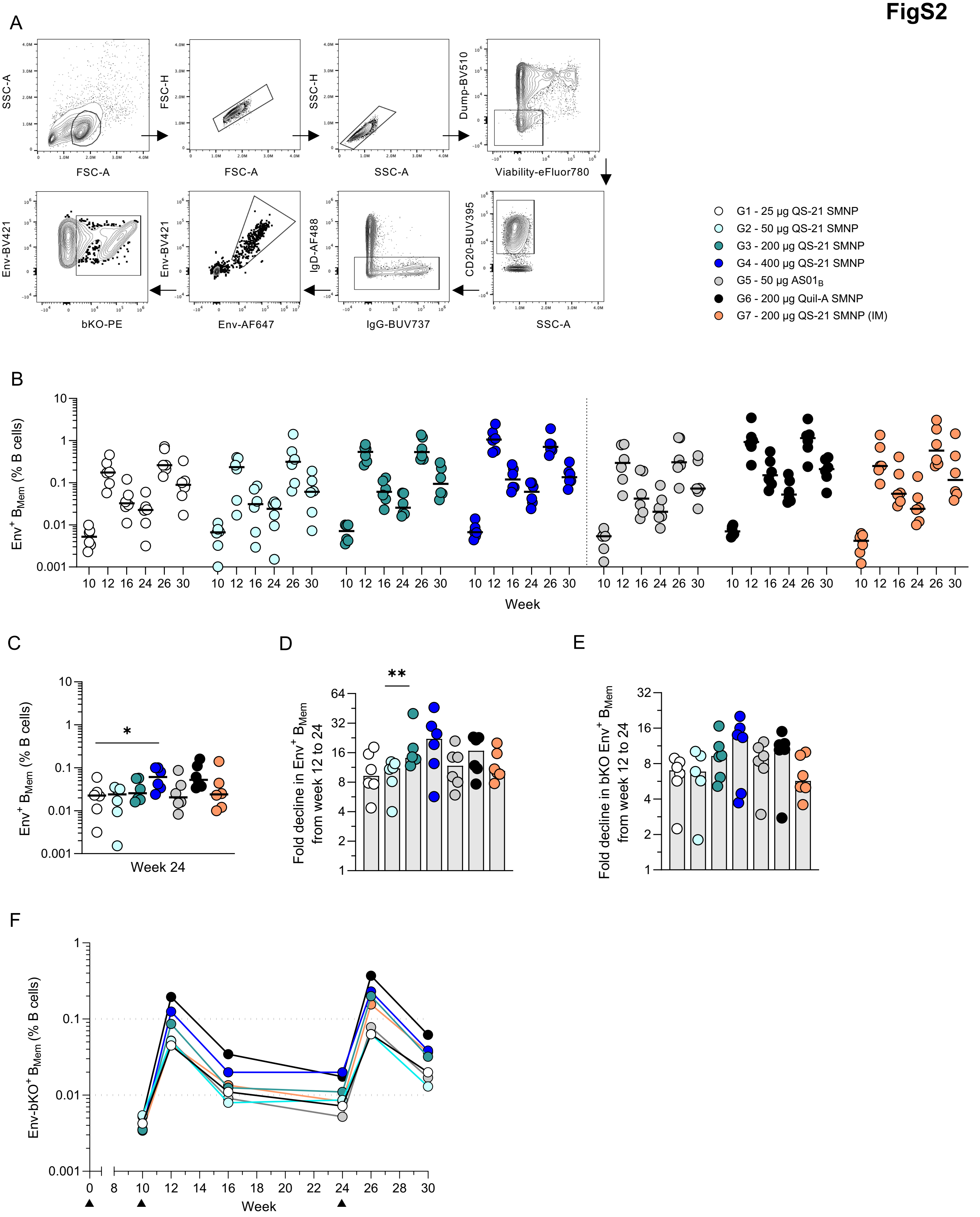
A
SSC-A
FSC-A
FSC-H
FSC-A
SSC-H
SSC-A
Dump-BV510
Viability-eFluor780
CD20-BUV395
SSC-A
IgD-AF488
IgG-BUV737
Env-BV421
Env-AF647
Env-BV421
bKO-PE
G1 - 25 µg QS-21 SMNP
G2 - 50 µg QS-21 SMNP
G3 - 200 µg QS-21 SMNP
G4 - 400 µg QS-21 SMNP
G5 - 50 µg AS01B
G6 - 200 µg Quil-A SMNP
G7 - 200 µg QS-21 SMNP (IM)
B
Env+ BMem (% B cells)
10 12 16 24 26 30
Week
C
Env+ BMem (% B cells)
*
Week 24
D
**
Fold decline in Env+ BMem from week 12 to 24
E
Fold decline in bKO Env+ BMem from week 12 to 24
F
Env-bKO+ BMem (% B cells)
0 8 10 12 14 16 18 20 22 24 26 28 30
Week

**Supplemental Figure 2. Gating strategy of Env-binding $B_{Mem}$ in PBMCs.**
**(A)** Flow cytometry gating strategy to define Env-binding $B_{Mem}$ in the blood.
**(B)** Frequency of Env-binding $B_{Mem}$ as a percentage of B cells.
**(C)** Frequency of Env-binding $B_{Mem}$ as a percentage of B cells in PBMCs at week 24.
**(D)** Fold change in Env-binding $B_{Mem}$ from week 12 to week 24.
**(E)** Fold change in bKO-binding $B_{Mem}$ from week 12 to week 24.
**(F)** Median frequency of bKO-binding $B_{Mem}$ as a percentage of B cells. Black triangles denote the time of immunization.
Horizontal black bars in **(B)** and gray bars in **(C) and (E)** represent the median.

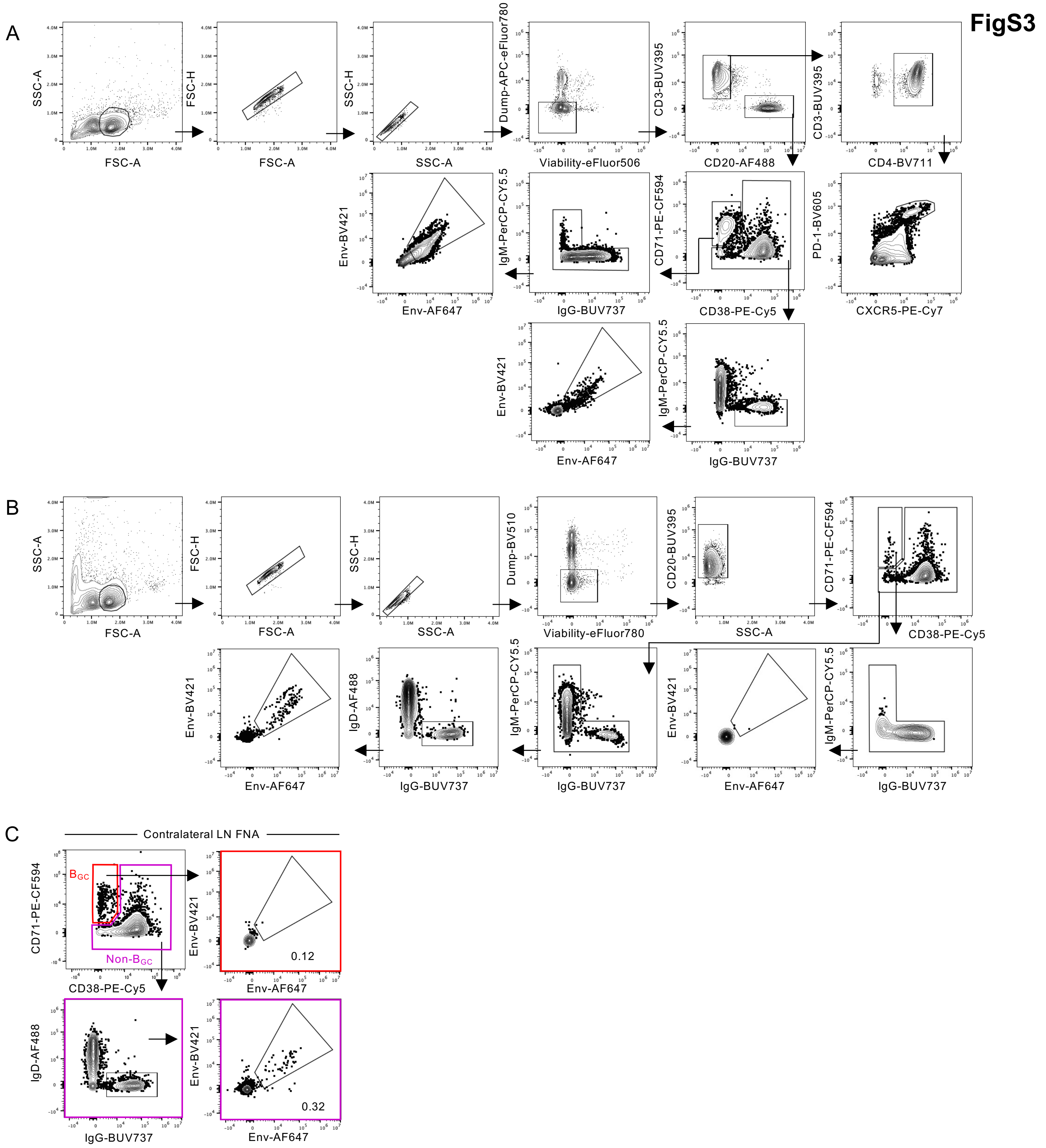
FigS3
A
SSC-A
FSC-A
FSC-H
FSC-A
SSC-H
SSC-A
Dump-APC-eFluor780
Viability-eFluor506
CD3-BUV395
CD20-AF488
CD3-BUV395
CD4-BV711
Env-BV421
Env-AF647
IgM-PerCP-CY5.5
IgG-BUV737
CD71-PE-CF594
CD38-PE-Cy5
PD-1-BV605
CXCR5-PE-Cy7
Env-BV421
Env-AF647
IgM-PerCP-CY5.5
IgG-BUV737
B
SSC-A
FSC-A
FSC-H
FSC-A
SSC-H
SSC-A
Dump-BV510
Viability-eFluor780
CD20-BUV395
SSC-A
CD71-PE-CF594
CD38-PE-Cy5
Env-BV421
Env-AF647
IgD-AF488
IgG-BUV737
IgM-PerCP-CY5.5
IgG-BUV737
Env-BV421
Env-AF647
IgM-PerCP-CY5.5
IgG-BUV737
C
Contralateral LN FNA
CD71-PE-CF594
CD38-PE-Cy5
$B_{GC}$
Non-$B_{GC}$
Env-BV421
Env-AF647
0.12
IgD-AF488
IgG-BUV737
Env-BV421
Env-AF647
0.32

**Supplemental Figure 3. Gating strategy of Env-binding $B_{GC}$ and $B_{Mem}$ in LN FNAs.**

**(A)** Flow cytometry gating strategy to define Env- binding $B_{GC}$ and $B_{Mem}$ in ipsilateral LN FNAs.

**(B)** Flow cytometry gating strategy to define Env- binding $B_{GC}$ and $B_{Mem}$ in contralateral LN FNAs.

**(C)** Flow cytometry gating of Env-binding $B_{GC}$ ($CD71^{+}CD38^{-}$) and $B_{Mem}$ (non-$B_{GC}$ $IgD^{-}IgG^{+}$) cells in contralateral LN FNAs at week 13.

**FigS4**

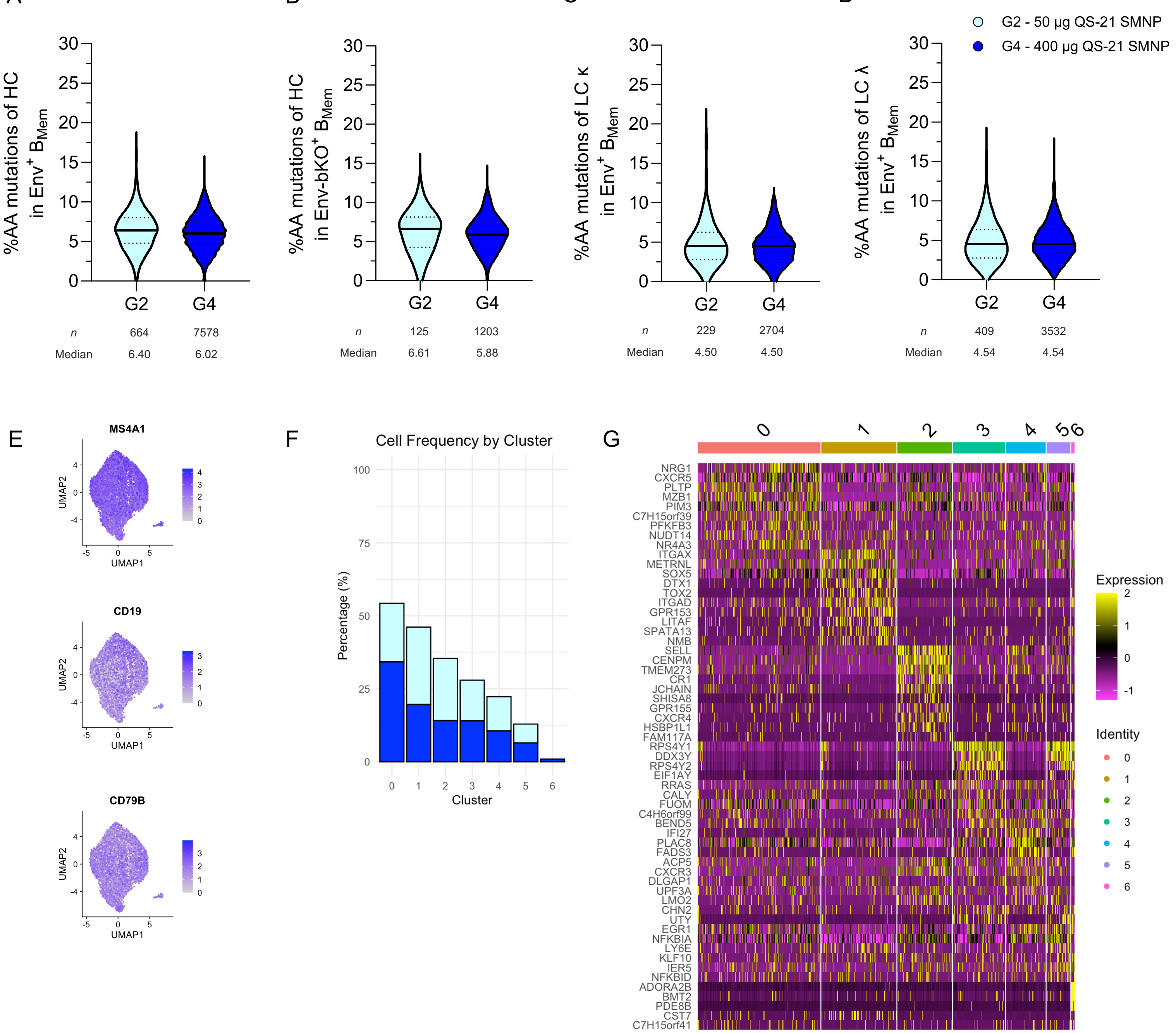

A
B
C
D
G2 - 50 µg QS-21 SMNP
G4 - 400 µg QS-21 SMNP
%AA mutations of HC in Env+ BMem
%AA mutations of HC in Env-bKO+ BMem
%AA mutations of LC κ in Env+ BMem
%AA mutations of LC λ in Env+ BMem
0
5
10
15
20
25
30
G2
G4
n 664 7578
Median 6.40 6.02
n 125 1203
Median 6.61 5.88
n 229 2704
Median 4.50 4.50
n 409 3532
Median 4.54 4.54
E
MS4A1
CD19
CD79B
UMAP1
UMAP2
F
Cell Frequency by Cluster
Percentage (%)
Cluster
0 1 2 3 4 5 6
0 25 50 75 100
G
NRG1
CXCR5
PLTP
MZB1
PIM3
C7H15orf39
PFKFB3
NUDT14
NR4A3
ITGAX
METRNL
SOX5
DTX1
TOX2
ITGAD
GPR153
LITAF
SPATA13
NMB
SELL
CENPM
TMEM273
CR1
JCHAIN
SHISA8
GPR155
CXCR4
HSBP1L1
FAM117A
RPS4Y1
DDX3Y
RPS4Y2
EIF1AY
RRAS
CALY
FUOM
C4H6orf99
BEND5
IFI27
PLAC8
FADS3
ACP5
CXCR3
DLGAP1
UPF3A
LMO2
CHN2
UTY
EGR1
NFKBIA
LY6E
KLF10
IER5
NFKBID
ADORA2B
BMT2
PDE8B
CST7
C7H15orf41
Expression
2
1
0
-1
Identity
0
1
2
3
4
5
6

**Supplemental Figure 4. Gene expression clusters of Env-binding $B_{Mem}$.**

**(A-B)** Percent of heavy chain (HC) amino acid (AA) mutations in **(A)** Env-binding $B_{Mem}$ and **(B)** bKO-binding $B_{Mem}$ cells at week 12 in PBMCs.

**(C-D)** Percent of light chain (LC) **(C)** kappa and LC **(D)** lambda amino acid (AA) mutations in Env-binding $B_{Mem}$ at week 12.

**(E)** Uniform manifold approximation and projection (UMAP) of single-cell gene expression profiles for *MS4A1*, *CD19* and *CD79B* in Env-binding $B_{Mem}$ at week 12.

**(F)** Cell frequency per cluster in Group 2 (50 µg QS-21 SMNP) and Group 4 (400 µg QS-21 SMNP) Env-binding $B_{Mem}$.

**(G)** Differential gene expression among clusters of $B_{Mem}$ at week 12.

The solid black line and dotted lines in **(A-D)** represent the median and quartiles, respectively. Statistical significance was determined by an unpaired two-tailed Mann-Whitney test. *$P \leq 0.05$, **$P < 0.01$, ***$P < 0.001$, ****$P < 0.0001$.

FigS5

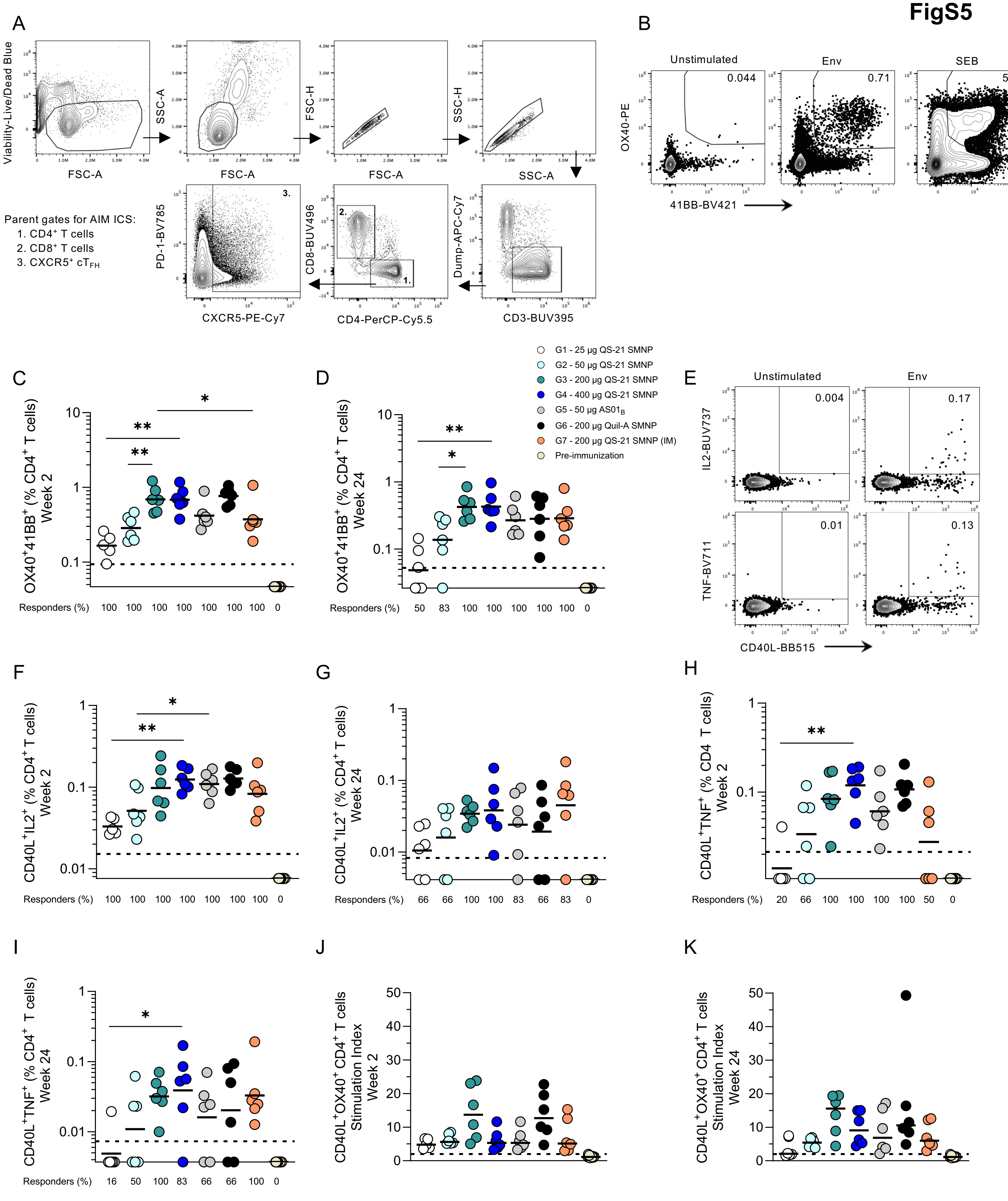

A
Viability-Live/Dead Blue
FSC-A
SSC-A
FSC-H
SSC-H
PD-1-BV785
CXCR5-PE-Cy7
CD8-BUV496
CD4-PerCP-Cy5.5
Dump-APC-Cy7
CD3-BUV395
Parent gates for AIM ICS:
1. CD4+ T cells
2. CD8+ T cells
3. CXCR5+ cTFH
B
Unstimulated
Env
SEB
0.044
0.71
5.83
OX40-PE
41BB-BV421
C
D
OX40+41BB+ (% CD4+ T cells) Week 2
OX40+41BB+ (% CD4+ T cells) Week 24
Responders (%) 100 100 100 100 100 100 100 0
Responders (%) 50 83 100 100 100 100 100 0
G1 - 25 µg QS-21 SMNP
G2 - 50 µg QS-21 SMNP
G3 - 200 µg QS-21 SMNP
G4 - 400 µg QS-21 SMNP
G5 - 50 µg AS01B
G6 - 200 µg Quil-A SMNP
G7 - 200 µg QS-21 SMNP (IM)
Pre-immunization
E
Unstimulated
Env
0.004
0.17
0.01
0.13
IL2-BUV737
TNF-BV711
CD40L-BB515
F
G
H
CD40L+IL2+ (% CD4+ T cells) Week 2
CD40L+IL2+ (% CD4+ T cells) Week 24
CD40L+TNF+ (% CD4 T cells) Week 2
Responders (%) 100 100 100 100 100 100 100 0
Responders (%) 66 66 100 100 83 66 83 0
Responders (%) 20 66 100 100 100 100 50 0
I
J
K
CD40L+TNF+ (% CD4+ T cells) Week 24
Responders (%) 16 50 100 83 66 66 100 0
CD40L+OX40+ CD4+ T cells Stimulation Index Week 2
CD40L+OX40+ CD4+ T cells Stimulation Index Week 24

**Supplemental Figure 5. Gating strategy of Env-specific CD4 and CD8 T cells.**
**(A)** Flow cytometry gating strategy to analyze Env-specific CD4 T cells, CD8 T cells and $cT_{FH}$ cells.
**(B)** Representative flow cytometry plots of $AIM^+$ $OX40^+41BB^+$ CD4 T cells from unstimulated (negative control) and following 24-hour stimulation with a BG505 MD39 Env peptide pool or superantigen staphylococcal enterotoxin (SEB, positive control) at week 2.
**(C-D)** Frequency of $AIM^+$ $OX40^+41BB^+$ Env-specific CD4 T cells at **(C)** week 2 and **(D)** week 24.
**(E)** Representative flow cytometry plots of $CD40L^+IL\text{-}2^+$ CD4 T cells and $CD40L^+TNF^+$ CD4 T cells from unstimulated (negative control) and following 24-hour stimulation with a BG505 MD39 Env peptide pool at week 2.
**(F-G)** Frequency of $CD40L^+IL\text{-}2^+$ Env-specific CD4 T cells at **(F)** week 2 and **(G)** week 24.
**(H-I)** Frequency of $CD40L^+TNF^+$ Env-specific CD4 T cells at **(H)** week 2 and **(I)** week 24.
**(J-K)** Stimulation index of $AIM^+$ ($OX40^+CD40L^+$) CD4 T cells at **(J)** week 2 and **(K)** week 24.
Horizonal black lines represent the geometric mean. Black dotted lines indicate the limit of quantification (LOQ). Percent responders was calculated as the percent of animals above the LOQ. Graphed are the frequencies after subtracting from paired unstimulated samples. Statistical significance was determined by an unpaired two-tailed Mann-Whitney test. *$P \leq 0.05$, **$P < 0.01$, ***$P < 0.001$, ****$P < 0.0001$.

FigS6

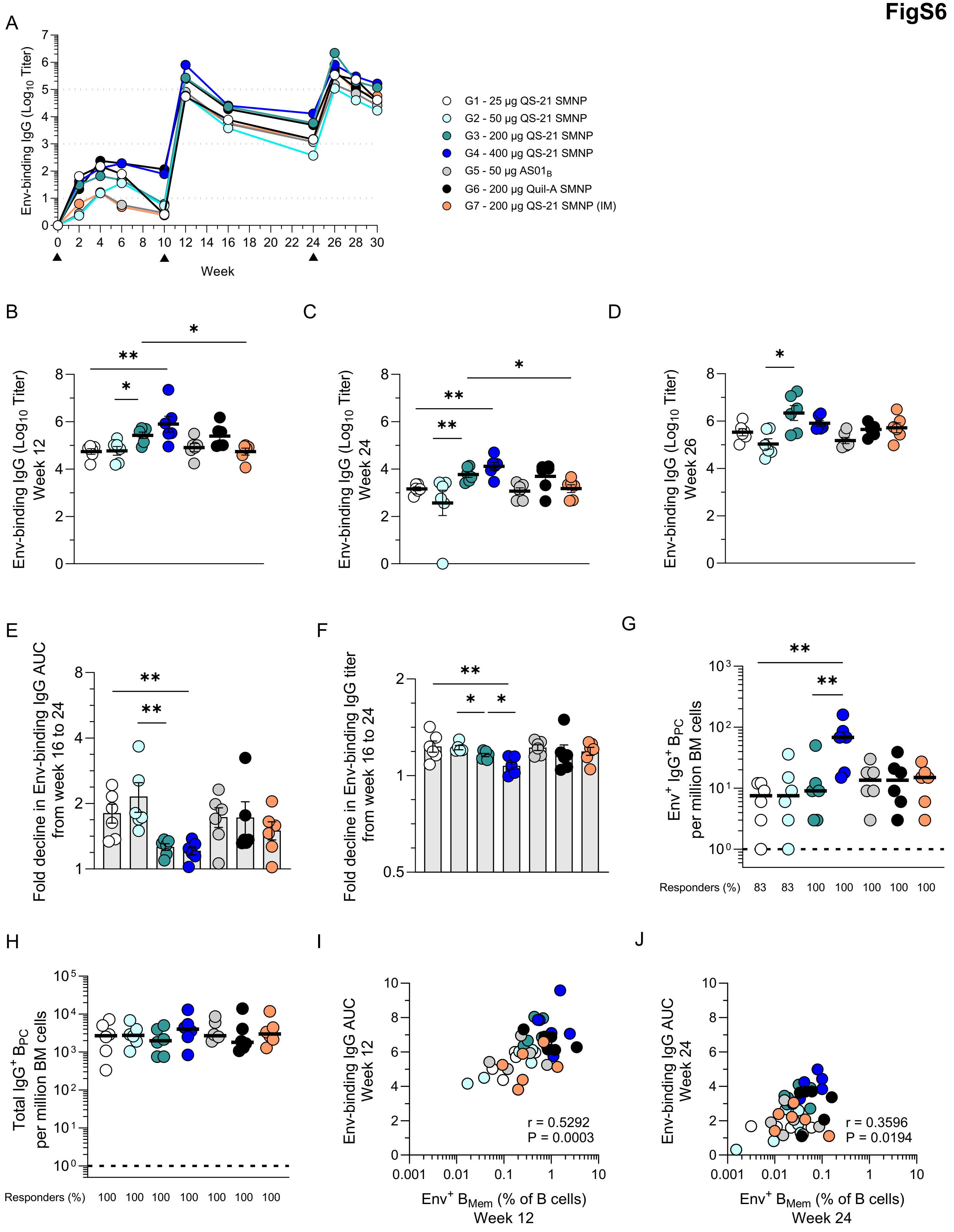
A
Env-binding IgG (Log10 Titer)
Week
G1 - 25 µg QS-21 SMNP
G2 - 50 µg QS-21 SMNP
G3 - 200 µg QS-21 SMNP
G4 - 400 µg QS-21 SMNP
G5 - 50 µg AS01B
G6 - 200 µg Quil-A SMNP
G7 - 200 µg QS-21 SMNP (IM)
B
Env-binding IgG (Log10 Titer) Week 12
C
Env-binding IgG (Log10 Titer) Week 24
D
Env-binding IgG (Log10 Titer) Week 26
E
Fold decline in Env-binding IgG AUC from week 16 to 24
F
Fold decline in Env-binding IgG titer from week 16 to 24
G
Env+ IgG+ BPC per million BM cells
Responders (%) 83 83 100 100 100 100 100
H
Total IgG+ BPC per million BM cells
Responders (%) 100 100 100 100 100 100 100
I
Env-binding IgG AUC Week 12
r = 0.5292
P = 0.0003
Env+ BMem (% of B cells) Week 12
J
Env-binding IgG AUC Week 24
r = 0.3596
P = 0.0194
Env+ BMem (% of B cells) Week 24

**Supplemental Figure 6. Env-binding IgG antibody titer responses.**
**(A)** Mean Env-binding IgG endpoint titers.
**(B-D)** Env-binding IgG endpoint titers at **(B)** week 12, **(C)** week 24 and **(D)** week 26.
**(E)** Fold change in Env-binding IgG AUC from week 16 to week 24.
**(F)** Fold change in Env-binding IgG endpoint titer from week 16 to week 24.
**(G)** Env-binding $B_{PC}$ ($IgG^{+}$) per million bone marrow cells at week 37.
**(H)** Total $B_{PC}$ ($IgG^{+}$) per million bone marrow cells at week 37.
**(I-J)** Correlation between Env-binding IgG antibodies and Env-binding $B_{Mem}$ at **(I)** week 12 and **(J)** week 24.
Error bars in **(B-F)** represent mean with SEM. Horizontal black bars in **(G-H)** represents the median. Dotted line in **(G-H)** denotes the limit of detection (LOD) and was used to calculate percent responders. For **(B-H)** statistical significance was determined by an unpaired two-tailed Mann-Whitney test. Data in **(I-J)** was analyzed using a Spearman's Correlation test. $*P \leq 0.05$, $**P < 0.01$, $***P < 0.001$, $****P < 0.0001$.

**Table S1. GMP-process SMNP characterization**

| Characteristic | Result |
|---|---|
| appearance | Colorless, slightly opalescent |
| Content QS-21 | 820 µg/mL |
| Content MPLA | 84 µg/mL |
| Content cholesterol | 196 µg/mL |
| Content DPPC | 93 µg/mL |
| Particle size (z-average) | 49 nm |
| Polydispersity index (PDI) | 0.15 |
| Residual MEGA-10 | < 1 µg/mL |
| pH | 6.5 |
| Osmolality | 281 mOsmol/kg |
| Endotoxin | < 1 EU/mL |
| Bioburden | 0 CFU / 10 mL |

**Table S2. Dilution steps for SMNP GMP-process synthesis**

| Dilution step | Dilution factor | Hold time between adding steps |
|---|---|---|
| 1 | 10x | 10 min |
| 2 | 25x | 45 min |
| 3 | 30x | 20 min |
| 4 | 35x | 20 min |
| 5 | 40x | 10 min |
| 6 | 45x | 10 min |
| 7 | 50x | 10 min |
| 8 | 60x | 10 min |
| 9 | 75x | 10 min |
| 10 | 85x | 10 min |
| 11 | 100x | 10 min |
